# Astrocytes amplify neurovascular coupling to sustained activation of neocortex in awake mice

**DOI:** 10.1101/2020.12.16.422785

**Authors:** Adam Institoris, Milène Vandal, Govind Peringod, Christy Catalano, Cam Ha Tran, Xinzhu Yu, Frank Visser, Cheryl Breiteneder, Leonardo Molina, Baljit S. Khakh, Minh Dang Nguyen, Roger J. Thompson, Grant R. Gordon

## Abstract

Brain requires increased local cerebral blood flow (CBF) for as long as necessary during neuronal activation to match O_2_ and glucose supply with demand – termed functional hyperemia. Ca^2+^ elevation in astrocytes can drive arteriole dilation to increase CBF, yet affirmative evidence for the necessity of astrocytes in functional hyperemia *in vivo* is lacking. In awake and active mice, we discovered that functional hyperemia is bimodal with a distinct early and late component whereby arteriole dilation progresses as sensory stimulation is sustained. Clamping astrocyte Ca^2+^ signaling *in vivo* by expressing a high-affinity plasma membrane Ca^2+^ ATPase (CalEx) reduces sustained but not brief sensory-evoked arteriole dilation. Reciprocally, elevating astrocyte free Ca^2+^ using chemogenetics selectively augments sustained but not brief hyperemia. Neither locomotion, arousal, nor changes in neuronal signaling account for the selective effect of astrocyte Ca^2+^ on the late phase of the CBF response. Antagonizing NMDA-receptors or epoxyeicosatrienoic acid production reduces only the late component of functional hyperemia, leaving brief increases in CBF to sensory stimulation intact. We propose that a fundamental role of astrocyte Ca^2+^ is to amplify functional hyperemia when neuronal activation is prolonged.

## Introduction

Neuronal function requires tight and uninterrupted access to O_2_ and glucose via the blood supply because neurons have neither sufficient energy stores nor capacity for anaerobic metabolism. Thus, neural processing of sensory, motor, and cognitive information drives increased regional cerebral blood flow (CBF) for seconds to minutes as needed. This phenomenon – functional hyperemia – is essential for brain metabolism and also serves as the basis of the blood oxygen level-dependent (BOLD) signal of functional magnetic resonance imaging (fMRI)^1,2^.

Functional hyperemia occurs, in part, by large diameter changes in penetrating arterioles ^3–6^ as a result of multiple parallel neurovascular coupling mechanisms thought to be governed by neurons, astrocytes and endothelial cells that actuate vascular smooth muscle. Astrocytes respond to neural activity and their peri-vascular processes/endfeet can release Ca^2+^ dependent messengers to regulate arteriole diameter, both *ex vivo* (reviewed in ^7,8^) and *in vivo* ^9,10^ when artificially stimulated. We have previously described that astrocytes sense behavioral-state and vascular signals as brief functional hyperemia evolves in awake mice ^11^, yet, their capacity to control CBF to a natural stimulus *in vivo* has not yet been demonstrated. It remains possible that astrocytes contribute to functional hyperemia only under distinct durations of neural activity. Indeed, astrocyte Ca^2+^ correlates to CBF when sensory stimulation is prolonged ^12^. Under anesthesia, astrocyte Ca^2+^ signals can be suppressed ^13^, and arteriole/CBF responses to sustained sensory stimulation consist of an initial rise followed by a plateau at close to the same amplitude ^14^. The latter has prompted many to solely explore initiating mechanisms or assume that there are no temporally distinct components to functional hyperemia. More recent studies, some conducted in awake animals, display CBF/arteriole response curves that appear bimodal ^15,16,17^, yet these temporally distinct phases are rarely mechanistically explored or even described. We hypothesized that delayed astrocyte Ca^2+^ signals (>3sec) occurring in the awake state regulate CBF when sensory activation is sustained, therefore comprising an important and unrecognized mechanism of functional hyperemia ^12,18^. Thus far, no causal ‘necessity’ and ‘sufficiency’ experiments support this hypothesis. The main framework of astrocyte Ca^2+^ in functional hyperemia *in vivo* relies on correlational studies ^12,18–23^, none of which have clearly elucidated the physiological context of astrocytic contributions at arterioles. Though recent work suggests a Ca^2+^ independent astrocytic mechanism in CBF regulation involving CO_2_^24^, the foundational work of this field manipulated and measured astrocyte free Ca^2+ 7,8^, highlighting the need to resolve the Ca^2+^ signaling hypothesis.

Using 2-photon fluorescence imaging in awake mice and tools to selectively clamp or augment astrocyte Ca^2+^ *in vivo*, we discovered that during sustained bouts of sensory activation 1) functional hyperemia increases bimodally and 2) astrocyte Ca^2+^ is causal for the augmentation of the second phase. These findings are important because impairments in sustained functional hyperemia occur in attention deficit ^25^, age-related cognitive impairment ^26^, dementia ^27^ and stroke ^28^. Our results help clarify neuro-glio-vascular coupling and will facilitate future work aimed at understanding how astrocytes impact the (patho)physiology of the cerebrovasculature.

## Results

### Sustained functional hyperemia escalates and is associated with astrocyte Ca^2+^ transients

We investigated the temporal dynamics of penetrating arteriole diameter and astrocyte Ca^2+^ during brief (5sec) and sustained (30sec) whisker stimulation using 2-photon microscopy through an acutely installed cranial window over the barrel cortex of awake mice (Fig. 1a). First, imaging *Aldh1l1*-Cre/ERT2 x RCL-GCaMP6s mice (N=9) arteriole responses to whisker stimulation with air puff elicited rapid arteriole dilation that peaked within the first 5sec of stimulation, followed by progressively increasing diameter only when stimulation was maintained for 30sec revealing a bimodal response (Fig. 1b,c). Interestingly, peak dilations to 1sec (Δd/d=6.07±1.2%) and 5sec stimulation (Δd/d=8.3±1.1%) were not different, but dilation to 30sec stimulation (Δd/d=25.95±5.7%) was significantly higher due to the putative secondary phase (Friedman-test P=0.001, Trial (T)=16-24, Penetrating Arteriole (PA)=7, N=6; Fig. 1d). Arteriole responses were similar in c57bl/6 mice imaged through thinned skull (t-test P=0.004, T= 17, PA=6, N=3; Suppl. Fig. 1a-c), confirming that acute craniotomy did not impair functional hyperemia. Astrocyte fine processes, followed by endfeet had Ca^2+^ signals that both increased after the onset of vasodilation (One way ANOVA P=0.0001, Fig. 1e,f)^11^. We found similar arteriole response dynamics and delayed elevation in Ca^2+^ within endfeet of the membrane-tethered GCaMP6 expressing *Aldh1l1*-Cre/ERT2 x R26-Lck-GCaMP6f mice implanted with a chronic cranial window over an intact dura (>4weeks recovery) (Suppl. Fig. 2a-f). We identified two distinct populations of fine process Ca^2+^ signals: 47% of signals started in the first sec of stimulation (ultrafast) before initiation of vasodilation and 45% appeared 3-5sec after stimulation onset (delayed)(Suppl. Fig. 2d,e) however, ultrafast signals ^19,21^ only reached a Ca^2+^ rise of ΔF/F=3.6±1.3% in the first sec (in contrast to ΔF/F= 22±3.5% at 3-5sec and ΔF/F=46.8±5.7% of the peak Ca^2+^ response, Friedman test P<0.0001, T=27) (Suppl. Fig. 2d). In further support of a bimodal arteriole response, imaging vascular smooth muscle cell Ca^2+^ changes in *PDGFRß*-Cre x RCL-GCaMP6s (T=30, PA=30, N=11) mice during 30sec whisker stimulation showed two separate peaks of Ca^2+^ drop at ∼2.5sec and between 20-30sec (Fig. 1g,h). This suggests separate processes mediate smooth muscle relaxation in the early and late phase of sustained functional hyperemia.

**Figure 1.**
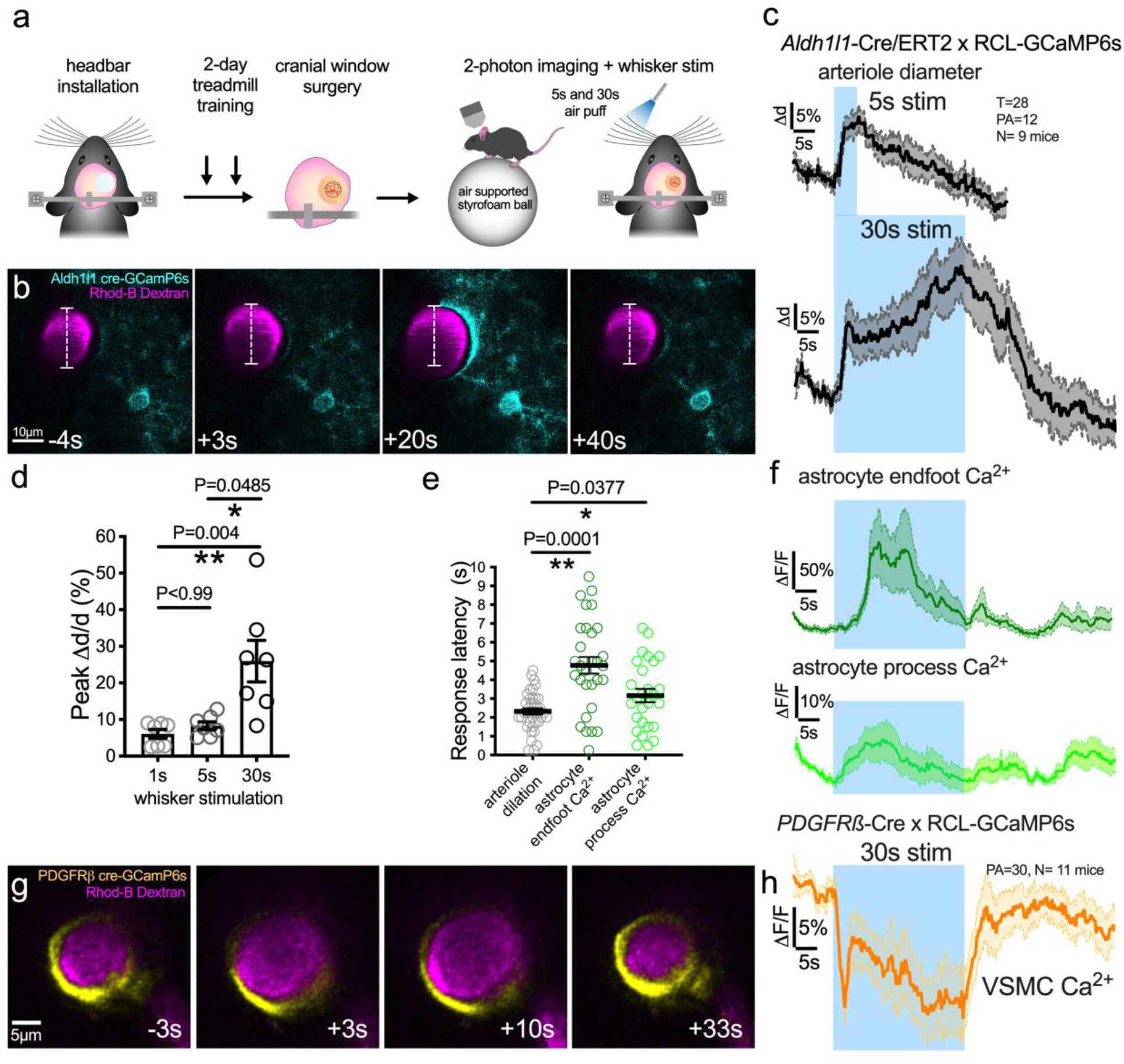
Sustained functional hyperemia escalates and is associated with delayed astrocyte Ca^2+^ transients. **a**) Cartoon and timeline of the acute cranial window with skull/dura removal preparation in awake mouse 2-photon imaging experiment in barrel cortex. **b**) Time series images showing dilation of a Rhodamine-B-dextran labelled penetrating arteriole and Ca^2+^ responses of a GCaMP6s-expressing astrocyte in response to whisker stimulation at 5sec and 30sec in an *Aldh1l1*-CreERT2 x RCL-GCaMP6s mouse. Time stamps refer to stimulation onset as 0sec. **c**) Traces of arteriole dilation (black) to 5 and 30sec whisker stimulation. **d**) Summary data of peak arteriole dilation to 1, 5 and 30sec stimulation. **e**) Summary of response onset (calculated from each trial as 3 x SD above baseline) for dilation and astrocyte Ca^2+^. **f**) Astrocyte endfoot (dark green) and astrocyte arbor (light green) Ca^2+^ traces to 30sec whisker stimulation. **g**) Time series images of GCaMP6s expressing vascular smooth muscle cells (VSMC)(yellow) within a penetrating arteriole wall loaded with Rhodamine-B-Dextran (magenta), and **h**) VSCM Ca^2+^ trace during 30sec whisker stimulation. Each image is an average projection of 4 raw images. All trace data show the average with SEM.

### Clamping astrocyte Ca^2+^ reduces the late phase of functional hyperemia

Given the stronger association between astrocyte process/endfoot Ca^2+^ and sustained functional hyperemia, and the appearance of a distinct late component in smooth muscle Ca^2+^ traces, we tested *ex vivo* and *in vivo* for a causal role of astrocyte Ca^2+^ primarily in the late phase of functional hyperemia. Astrocyte Ca^2+^ transients mediate activity-dependent capillary but not arteriole dilation to 5sec stimulation in acute cortical brain slices ^29^. Thus, we tested for a causal relationship between long (30sec) but not short (5sec) stimulation-evoked astrocyte Ca^2+^ and penetrating arteriole dilation in acute rat brain slices, by whole-cell patch infusing the Ca^2+^ chelator BAPTA (10mM) into a network of peri-arteriolar astrocytes. We clamped intracellular free Ca^2+^ close to the resting concentration in astrocytes (∼100nM) ^30,31^. We allowed 15 mins for the internal solution to equilibrate in the astrocyte network ^32^ and for arteriole tone to stabilize. Rhod-2/AM was used to label large astrocyte and neuronal compartments, as well as the neuropil. Before patching an astrocyte, 30sec of high frequency electrical stimulation elicited Ca^2+^ transients in neurons, the neuropil, astrocyte somata and endfeet, as well as caused arteriole dilation (N=8) (Suppl. Fig. 3b-e). This was then repeated when the astrocyte network was filled with either control internal or Ca^2+^ clamp internal solution. We found that only the Ca^2+^ clamp solution reduced arteriole dilation to 30sec afferent stimulation (unpaired t test, N=5-8, P=0.039). While the astrocyte network filling itself caused reductions in evoked Ca^2+^ from all cell compartments, comparing to the Ca^2+^ clamp solution revealed that the only effect to explain the loss of dilation was the clamp on astrocyte Ca^2+^ (Mann-Whitney test, P=0.029, N=5-8, Suppl. Fig. 3d-e). Importantly, neither clamping astrocyte Ca^2+^, nor the control patch, affected arteriole dilation to 5sec stimulation, and patching itself (in any condition) did not affect baseline arteriole tone after whole-cell equilibration (Suppl. Fig. 4). These data show that longer neural stimulation is required to recruit Ca^2+^ dependent astrocyte contributions to arteriole dilation in neocortical brain slices.

Previous attempts to connect astrocyte Ca^2+^ to functional hyperemia in vivo focused on IP3R2 knockout ^33–35^, but to our knowledge there have been no *in vivo* manipulations to clamp astrocyte Ca^2+^ more generally for CBF control; thereby affecting multiple Ca^2+^ dependent pathways. We made use of the astrocyte Ca^2+^ silencing tool CalEx ^36^. Here, astrocyte-selective AAV (AAV2/5-gfaABC_1_D-CalEx-HA) delivers a high-affinity Ca^2+^ pump to the plasma membrane (Fig. 2a,b), which reduces evoked and spontaneous elevations in free Ca^2+^ both *in vitro* and *in vivo* ^36^. As a control, we made a single point mutation in the CalEx sequence (E457A) to prevent Ca^2+^ transport across the cell membrane ^37,38^. We validated both tools in awake c57bl/6 mice via intracortical viral injection followed by a chronic cranial window implantation (Fig. 2c). First, we compared astrocyte AAVs for CalEx and GCaMP6f vs the mutated CalEx control virus (AAV2/5-gfaABC_1_D-CalExMut(E457A)-HA) + AAV2/5-gfaABC_1_D-GCaMP6f), by examining startle evoked astrocyte Ca^2+^ signals in the barrel cortex (untrained body air puff, Supp. Fig. 5a,b). Astrocytes expressing functional CalEx showed markedly reduced startle-evoked Ca^2+^ transients (ΔF/F max=22.3±7.2%; AUC=277±105 arbitrary unit (a.u.), N=6) compared to control (ΔF/F max=90.9±25.8%, unpaired t test: P=0.0268; AUC=780±164 a.u., unpaired t test: P=0.0271, N=6, Suppl. Fig. 5c,d). *Post hoc* CalEx expression was confirmed by immunostaining against the reporter hemagglutinin (HA) (Fig. 2d).

**Figure 2.**
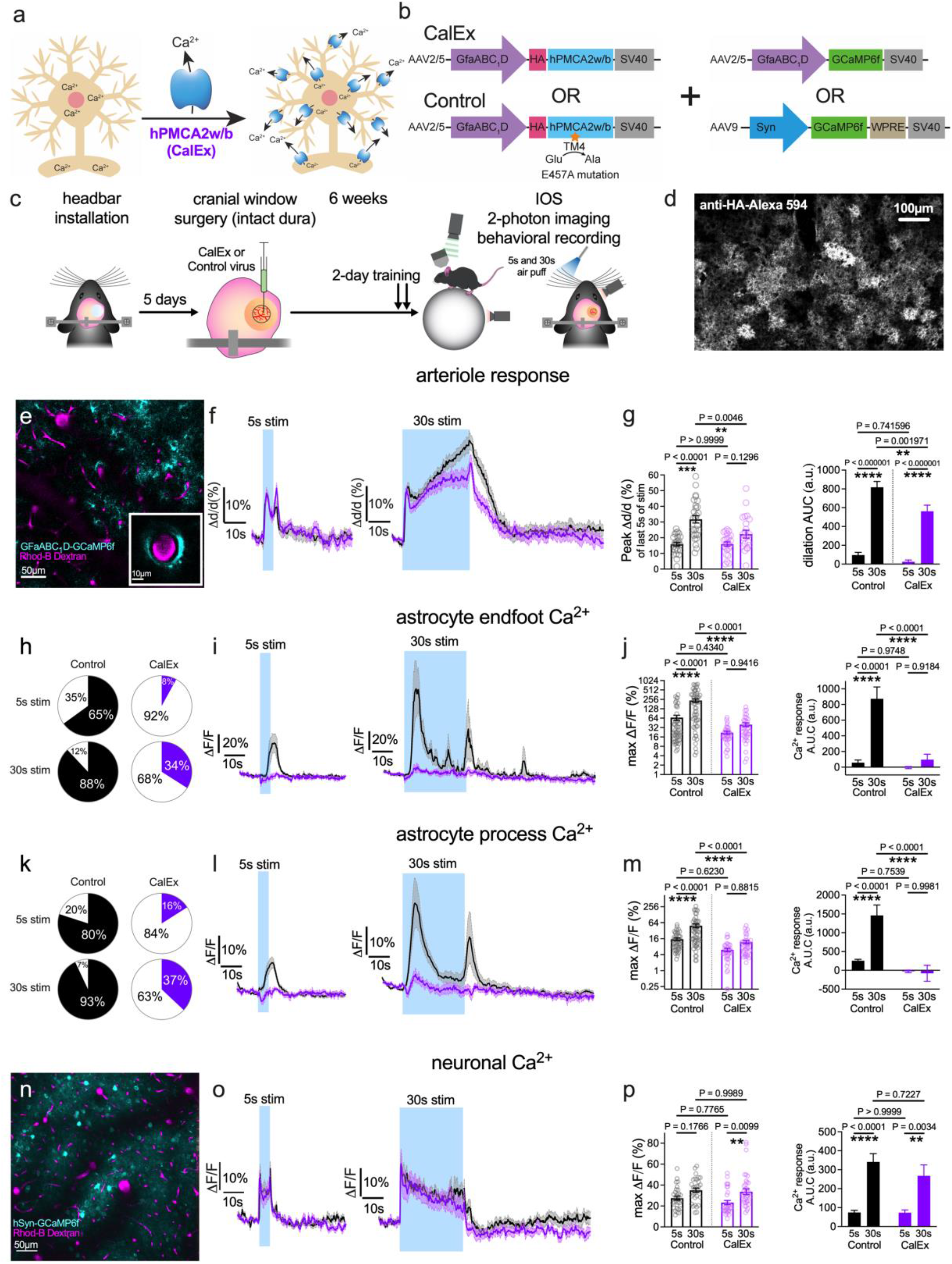
Clamping astrocyte Ca^2+^ reduces the late phase of functional hyperemia. **a**) Cartoon of astrocyte Ca^2+^ extrusion tool: a high-affinity plasma membrane Ca^2+^ ATPase hPMCA2w/b (CalEx). **b**) Viral vector strategy to express CalEx (*Top Left*) in astrocyte (or control virus-*Bottom Left*) plus GCaMP6f targeting astrocytes (*Top Right*) or neurons (*Bottom Right*). **c**) Timeline from viral vector intracortical injection to the imaging experiment. **d**) *Post hoc* immunofluorescence of CalEx expression against the fused Hemagglutinin (HA) reporter. **e**) Representative image of gfaABC_1_D-GCaMP6f expressing astrocytes (cyan) and Rhodamine-B-Dextran loaded vasculature (magenta) in layer 2/3 of the barrel cortex 6 weeks after viral injection. **f**) Average arteriole diameter traces with SEM in CalEx and control for 5sec (*Left*) and 30sec (*Right*) whisker stimulation. **g**) *Left:* summary data showing peak dilation of the last 5sec of the stimulation period and *Right:* area under the curve (AUC: 40sec from stimulation onset) in the four conditions. **h**) Astrocyte endfoot Ca^2+^ evoked event occurrence. Black (control) and purple (CalEx) slices are events, white is no event detected. An event is >3 standard deviation of baseline. **i**) Summary time series data of astrocyte endfoot Ca^2+^ in CalEx vs control for both 5sec (*Left*) and 30sec (*Right*) whisker stimulation. **j**) Summary of peak (*Left*) and AUC (stimulation+10sec) (*Right*) data in the four conditions. **k-m**) Same as *h-j* but for astrocyte arbor Ca^2+^. **n**) Representative image of AAV9.hSyn.GCaMP6f expressing neurons (cyan) and Rhodamine-B-Dextran loaded vasculature (magenta) in layer 2/3 of the barrel cortex 6 weeks after viral injection. **o**) Average neuronal Ca^2+^ traces with SEM in CalEx and control for 5sec (*Left*) and 30sec (*Right*) whisker stimulation. **p**) *Left:* summary data showing peak neuronal Ca^2+^ during the stimulation period and *Right:* area under the curve (AUC) of the stimulation period in the four conditions.

In response to 5sec whisker stimulation, we found that astrocyte CalEx had no impact on evoked arteriole dilation (peak Δd/d:16.0±1.4%) compared to control AAV (peak Δd/d: 15.9±1.1%, Tukey’s test: P=0.999, Fig. 2g). Yet remarkably, astrocytic CalEx reduced only the late component (last 5sec of stim) of arteriole dilation to 30sec whisker stimulation (peak Δd/d: 22.3±2.5%, T=68) compared to control (peak Δd/d: 31.7±2.3%, T=79, Fig. 2g) (Two-way ANOVA, P_CalEx_=0.0188, Tukey’s test: P=0.0046; CalEx: T=73, PA=23, N=10; control: T=85, PA=28, N=10; mean±SEM) (see Fig. 2g for stim+10s AUC data). These results could not be explained by differences in baseline arteriole tone (unpaired t test: P=0.369, N=27-23) caused by CalEx expression (Suppl. Fig. 6), and instead suggest that astrocyte Ca^2+^ is important for amplifying arteriole dilation during sustained, but not brief, functional hyperemia.

Confirming a clamp on astrocyte Ca^2+^, CalEx reduced the delayed astrocyte endfoot Ca^2+^ signal occurrence (Fig.2h) and peak caused by 5sec (Fig. 2i-j) and 30sec whisker stimulation (39.8±5.4% (T=38, PA=12, N=10) vs. 236±29.6% (T=57, PA=18, N=11), mean±SEM, Tukey’s test: P<0.0001), as well as the integrated Ca^2+^ response (AUC of stimulation period+10sec:) with an overall significance of P_CalEx_<0.0001 (Two-way ANOVA, Fig. 2j). CalEx also diminished astrocyte process Ca^2+^ transient occurrence (Fig.2k), Ca^2+^ peaks and Ca^2+^ response curves (AUC) when stimulating for 5sec (Fig. 2l) and for 30sec (12±1.8% (T=37) vs 49.2±7.8% (T=57), Two-way ANOVA, P_CalEx_<0.0001, Fig. 2l-m) (see Fig. 2m for AUC data). These data demonstrate that CalEx is an effective tool for clamping astrocyte Ca^*2+*^ signals *in vivo*.

It is well established that astrocyte Ca^2+^ signals and subsequent gliotransmission can shape synaptic transmission and neuronal excitability ^23,39–41^. For example, astrocyte-targeted CalEx in the striatum caused a loss of tonic inhibition in medium spiny inhibitory neurons, yet excitatory inputs onto these cells were unaltered ^23^. This prompted us to explore the effect of astrocyte CalEx on neuronal Ca^2+^ signals in the barrel cortex during brief and prolonged whisker stimulation by co-injecting CalEx or its control virus with a pan-neuronal GCaMP6f encoding virus (AAV9.hSyn.GCaMP6f). We recorded bulk neuronal Ca^2+^ signals over a set 5000μm^2^ area in layer2/3 around a responsive penetrating arteriole (Fig. 2n) and observed that neuronal Ca^2+^ peaked immediately after stimulation onset and gradually declined during 30s stimulation in both control and CalEx mice (Fig. 2o). This showed that the overall neural Ca^2+^ response (max ΔF/F and AUC of stim+10sec) was unaffected by attenuating astrocyte Ca^2+^ with CalEx (max ΔF/F: P_CalEx_ =0.242; AUC of stim+10sec: P_CalEx_ =0.5385, T=30-32; Fig. 2p). We also performed a more elaborate analysis of discrete Ca^2+^ transients occurring in various neuronal compartments in our field of view using an event-based toolkit ^42^. We calculated absolute event frequency, peak Ca^2+^ responses, area under the curve (AUC), absolute spatial area and duration of single events (control: 28905 events, T=56, PA=10, N=5 mice; CalEx: 39079 events, T=68, PA=11, N=5 mice, Suppl. Fig. 7). Some baseline differences between the groups were noted, which may be due to either a direct effect on local baseline neuronal activity or small differences in locomotor activity during this time (Suppl. Fig. 7g). The strongest effect of astrocyte CalEx on evoked neuronal calcium transients was the reduction in peak amplitudes, which was seen in both 5sec and 30sec stimulations (Suppl. Fig. 7b). However, both groups showed no effect on the initial vasodilation early in the response (see Fig. 2f), suggesting this change had no impact on early functional hyperemia. Additionally, during 30sec whisker stimulation, neuronal Ca^2+^ transients exhibited a small increase in AUC (Suppl. Fig. 7c), yet both the AUC and peak effects on neuronal Ca^2+^ signals were observed uniformly across the entire 30sec stimulation period – not selectively on the late phase. Collectively, these data exploring an indirect role for neuronal activity, did not adequately explain the preferential effect of astrocyte CalEx on the late component of functional hyperemia.

Cortical astrocytes integrate local sensory information and behavioral state ^11,43^. Locomotion, alone ^44,45^ or concomitantly with tactile stimulation of the whiskers, was shown to trigger astrocyte Ca^2+^ events ^46,44^, as well as to modulate cerebral blood flow in the neocortex ^17,47^. Notably, two-thirds of barrel cortex excitatory neurons respond predominantly to the combination of vibrissa deflection and running ^48^. Therefore, we assessed whether differences in relative locomotion during the stimulation period were confounding the differences in arteriole response and astrocyte Ca^2+^ between CalEx and control mice. Whisker stimulation prompted running at the onset and offset of whisker stimulation (Fig. 3a-b). Mice ran more during and after stimulation than at baseline. We found no difference in average locomotion during the 5sec (CalEx: 0.3127±0.01 vs control: 0.3024±0.01 a.u., Mann-Whitney test: P=0.1991) and 30sec (CalEx: 0.3181±0.01 vs control: 0.3247±0.01 a.u., Mann-Whitney test: P=0.8977) stimulation period between CalEx (T=69) and the control (T=75) group. Importantly, arteriole dilation and astrocyte Ca^2+^ signals returned to baseline after the stimulus (Fig 2f,i,l). This was despite mice running more after the stimulus than during the pre-stimulus period, showing that running per se did not drive a prolonged increase in arteriole diameter or astrocyte calcium elevation in the barrel. Co-variate analysis of locomotion during the stimulation period in a general linear model confirmed that running does not account for differences in arteriole (peak Δd/d: P=0.489, AUC: P=0.813) and neuronal Ca^2+^ (max ΔF/F: P=0.559, AUC: P=0.912) responses, as well as in astrocyte process Ca^2+^ (max ΔF/F: P=0.16, AUC: P=0.627) caused by CalEx. Interestingly, locomotion influenced the size of endfoot Ca^2+^ signals (max ΔF/F: P=0.001, AUC: P=0.02), but it did not negate the effect of CalEx on endfoot Ca^2+^ transients (P_CalEx_<0.0001). Collectively, these results suggest that differences in locomotion are unlikely to explain the differences in astrocyte Ca^2+^ and arteriole responses in the late phase of functional hyperemia.

**Figure 3.**
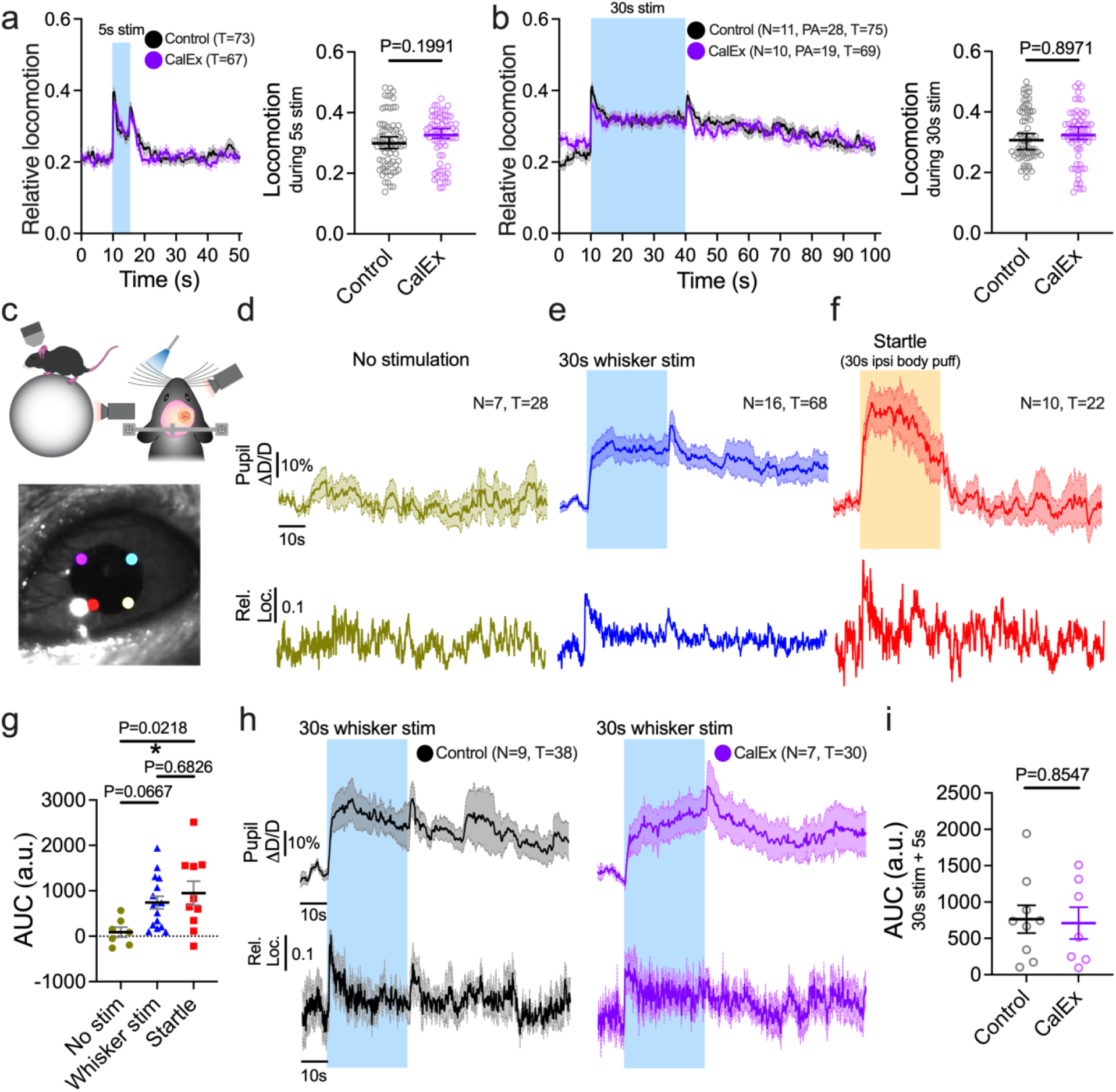
Locomotion and pupil size changes do not underlie CalEx effect during brief (5sec) and sustained (30sec) functional hyperemia. **a**) Summary curves (*Left*) and average values (*Right*) of relative locomotion during 5sec whisker stimulation for control and CalEx mice. **b**) Same but for 30sec whisker stimulation (mean ± SEM). **c**) Cartoon of locomotion (*Top Left*) and pupil (*Top Right*) recording with infrared cameras. *Bottom*: Pupil diameter tracking in two axes with the DeepLabCut tracking toolbox. **d**) Summary of relative pupil diameter changes (mean ± SEM) and relative locomotion (mean) during no stimulation recording, **e**) during 30sec whisker stimulation and **f**) during startle evoked by 30sec ipsilateral neck air puff (CalEx and Control combined). **g**) Averaged net Area Under the Curve (AUC) of pupil diameter for no stimulation, 30sec whisker stimulation + 5sec and 30sec startle + 5sec periods (mean ± SEM). **h**) Summary of relative pupil diameter changes (mean ± SEM) and relative locomotion (mean ± SEM) during 30sec whisker stimulation in control (*Left*) and CalEx mice (*Right*). **i**) Averaged net AUC of relative pupil diameter changes (stimulation + 5sec) in the control and CalEx groups.

Arousal provides a robust non-sensory modulation of brain activity across several sensory areas ^43,49,50^ and has been linked to astrocyte Ca^2+^ transients. The measurement of pupil diameter allows a good estimation of arousal state ^51^. We assessed and compared 30sec whisker stimulation-induced pupil diameter changes to a stimulus-free period and to a startle response elicited by 30sec ipsilateral air puff to the neck in a separate set of trials (Fig. 3c-i). To establish a bidirectionally responsive mid-size pupil, we used ambient green light. In trials with no stimulation, subtle changes in pupil diameter around the baseline corresponded to simultaneous changes in locomotion (AUC: 89±110 a.u., N=7) ^52^ (Fig. 3c,d). Whisker stimulation for 30sec showed a trend towards sustained pupil dilation (AUC: 740±140 a.u., One-way ANOVA and Tukey’s test: P=0.0667, N=16) that was at least partly reflected by an increase in locomotion (Fig. 3e,g). In contrast, the pupil dilated more robustly in response to startle (AUC: 948±262 a.u., P=0.0218, N=10) which was uncoupled from locomotion passed the initial onset period (Fig. 3f,g). These results indicate potentially heightened arousal to whisker air puff which was not as pronounced as a startle reaction. Finally, we confirmed that the pupil reaction in the CalEx group was not different from control (Unpaired t-test: P=0.8547, N=7-9), indicating a similar influence of arousal on functional hyperemia.

### Chemogenetic activation of astrocytes enhances only the late phase of functional hyperemia

Large astrocyte Ca^2+^ signals occur when synaptic glutamate or neuromodulators stimulate Gq-coupled receptors through inositol triphosphate (IP3) signaling ^22,53,54^. However, functional hyperemia is intact in anesthetized, sedated and awake mice lacking IP3-receptor 2 ^33–35,55,56^, the predominant subtype in astrocytes. Nevertheless, Gq-chemogenetics permits manipulation of cytosolic astrocyte Ca^2+^, which likely impacts signaling cascades outside of IP3R2 as a non-native receptor, allowing us to explore the contribution of general Gq signaling in awake mice. We aimed to test if 1) Gq-DREADD activation in astrocytes directly affects arteriole diameter, and if 2) continuous activation of astrocytic Gq either occludes or amplifies functional hyperemia to short vs long whisker stimulation. We crossed *Aldh1l1*-Cre/ERT2 with CAG-LSL-Gq-DREADD (N=6) mice to selectively express hM3Dq receptors on astrocytes, which was confirmed by visualizing the fusion reporter mCitrine (Fig. 4a). A perforated coverglass was applied to the brain surface to permit stable imaging while allowing topical superfusion of the DREADD agonist compound 21 (C21, 10μM). C21 triggered a prolonged (20-40min) astrocyte soma Ca^2+^ (ΔF/F=63±7%, paired t test: P<0.0001, N=14) and endfoot Ca^2+^ elevation (ΔF/F=34±4%, paired t test: P=0.0002, N=14), accompanied by transient arteriole dilation (Δd/d=19±2%, Wilcoxon test: P<0.0001, PA=15)(Fig. 4b-c). After an hour of continuous C21, astrocyte Ca^2+^ levels and arteriole diameter returned to baseline (Fig. 4d). Notably, arteriole dilation to 5sec whisker stimulation remained unchanged in the presence of C21 (peak Δd/d=12.2±1%) compared to pre-drug control (peak Δd/d=10.2±1%, paired t test: P=0.0533, PA=16, Fig. 4e), whereas 30sec stimulation showed a significant augmentation in only the later phase of functional hyperemia in C21 (Δd/d=24.7±2.3%), relative to before drug (Δd/d=17.5±1.9%, paired t test: P=0.0056, PA=16, N=6 mice, Fig. 4f) (See Fig. 4e,f for AUC data). Due to unexplained poor loading of Gq-DREADD positive astrocytes with Rhod2/AM, we could not adequately capture endfoot Ca^2+^ signals in response to whisker stimulation before and during C21 application. To test whether this effect is indirectly mediated via boosting neuronal activity, another group of *Aldh1l1*-Cre/ERT2 x CAG-LSL-Gq-DREADD mice received an AAV9.hSyn.GCaMP6f injection in the barrel cortex 4 weeks prior to imaging neuronal Ca^2+^ responses (Fig. 4g). Topical superfusion of C21 for 30min did not change bulk neuronal GCaMP6 signals at rest (ΔF/F=+5.4±0.17%, N=5, Fig. 4h) and failed to facilitate neuronal Ca^2+^ responses during 30sec whisker stimulation (Wilcoxon test: P=0.375, N=7, Fig. 4i). Our results demonstrate that selectively driving the astrocyte Gq-pathway causes arteriole dilation and only influences the late phase of sustained functional hyperemia.

**Figure 4.**
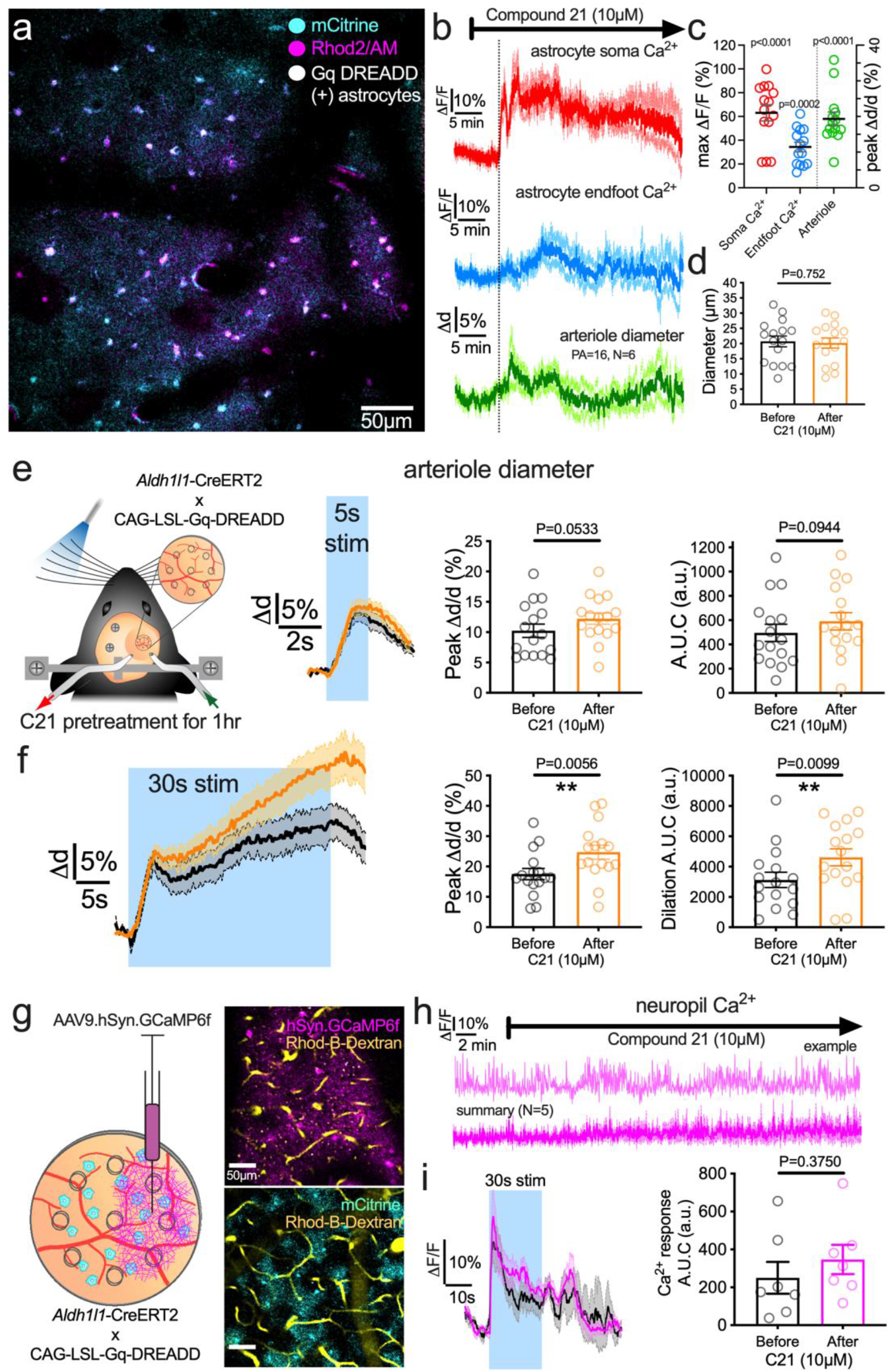
Chemogenetic activation of astrocytes enhances only the late phase of functional hyperemia. **a**) 2-photon image in barrel cortex of a *Aldh1l1*-Cre x CAG-LSL-Gq-DREADD mouse. Astrocytes are loaded with Rhod-2/AM and express Gq-DREADD-mCitrine. **b**) Time series measurements of astrocyte Ca^2+^ and arteriole diameter in response to the local superfusion of the DREADD agonist C21 into the perforated window. **c**) Summary data of peak responses in *b*. **d**) Summary data showing that in the continual presence of C21 arteriole diameter returns to baseline after 1hr. **e**) *Left*: Cartoon of awake mouse sensory stimulation experiment in the presence of C21 delivered via a perforated window. *Middle*: Average arteriole diameter traces with SEM in pre-drug control (black) and in the presence of C21 (orange), in response to 5sec whisker stimulation. *Right*: summary data showing peak dilation and AUC for 5sec in control and C21. **f**) Same as *e* but for 30sec whisker stimulation. **g**) *Left*: Cartoon showing the site of AAV9.Syn.GCaMP6f injection 4 weeks prior to acute cranial window experiment. *Top Right*: image of neuronal AAV9.hSyn.GCaMP6f expression (magenta) and Rhodamine-B-Dextran (yellow) loading of the vasculature in the lateral side of the window imaged at 920nm. *Bottom Right*: image of astrocytic mCitrine expression (cyan) in the medial side of the window imaged at 980nm. **h**) Time series measurements of neuronal Ca^2+^ in response to the local superfusion of the DREADD agonist C21 into the perforated window. Upper trace shows a representative ROI. Lower trace shows the averaged data. **i**) Average neuronal Ca^2+^ traces with SEM in pre-drug control (black) and in the presence of C21 (magenta), in response to 30sec whisker stimulation. *Right*: summary data showing AUC for 30sec stimulation before and after C21.

### NMDA receptors and EETs are involved in sustained functional hyperemia

N-methyl-D-aspartate receptors (NMDAR) are key participants in functional hyperemia. In the canonical model, synapses generate NMDAR-dependent nitric oxide, which causes rapid vasodilation (reviewed in ^57–59^). More recent work implicates NMDAR on astrocytes ^60^ and endothelium ^61^ in CBF control. Little is known about the contribution of NMDARs to different durations of functional hyperemia in awake animals. As NMDARs localize to the cell surface, we used *Aldh1l1*-Cre/ERT2 x R26-Lck-GCaMP6f mice (Fig. 5a) to express a plasma membrane-tethered GCaMP in astrocytes and imaged endfeet and processes surrounding a penetrating arteriole (Fig. 5c) or surrounding a precapillary sphincter – a control point for capillary access to blood ^4^. We again applied short (5sec) and long (30sec) duration whisker stimulation before and during topical NMDAR antagonist D,L-AP5 (1mM) via a perforated coverglass window. The lck-GCaMP6f showed similar astrocyte Ca^2+^ responses to the cytosolic GCaMP6s under control conditions (see Fig. 1f). Notably, in response to 5sec and 30sec whisker stimulation, D,L-AP5 eliminated both astrocyte endfoot (Two-way ANOVA: P_AP5_=0.0026, T=18-25, PA=7-8, N= 7-8 mice) and process (Two-way ANOVA: P_AP5_=0.0029) Ca^2+^ transients around penetrators (Fig. 5b,d), and a similar trend was observed in astrocytes around sphincters but failed to reach significance (Two-way ANOVA: P=0.0724, T=13-16, N=5-6, Suppl. Fig. 8). In a separate set of experiments using c57bl/6 mice (Fig. 5e), arteriole dilation to 5sec whisker stimulation remained surprisingly unchanged in the presence of D,L-AP5 (Δd/d=15.3±2.8%) from control (Δd/d=18.3±3.4%, Two-way ANOVA and Tukey’s test: P=0.9251, PA=8, N=7, Fig. 5f). Before NMDAR antagonism, vasodilation was again larger during 30sec stimulation (Δd/d=27±5.1%) compared to 5sec (Tukey’s test: P_Δd/d_=0.31, P_AUC_=0.0006, Fig. 5f), but in the presence of D,L-AP5, the late component of functional hyperemia was completely blocked, as arteriole diameter returned to baseline quickly after the initial peak (Δd/d=11.8±1.6%, Tukey’s test: P=0.0214; PA=8, N=7 mice, Fig. 5f) (see Fig. 5f for AUC data). These data show NMDARs are critical for astrocyte activation and for the late phase of functional hyperemia.

**Figure 5.**
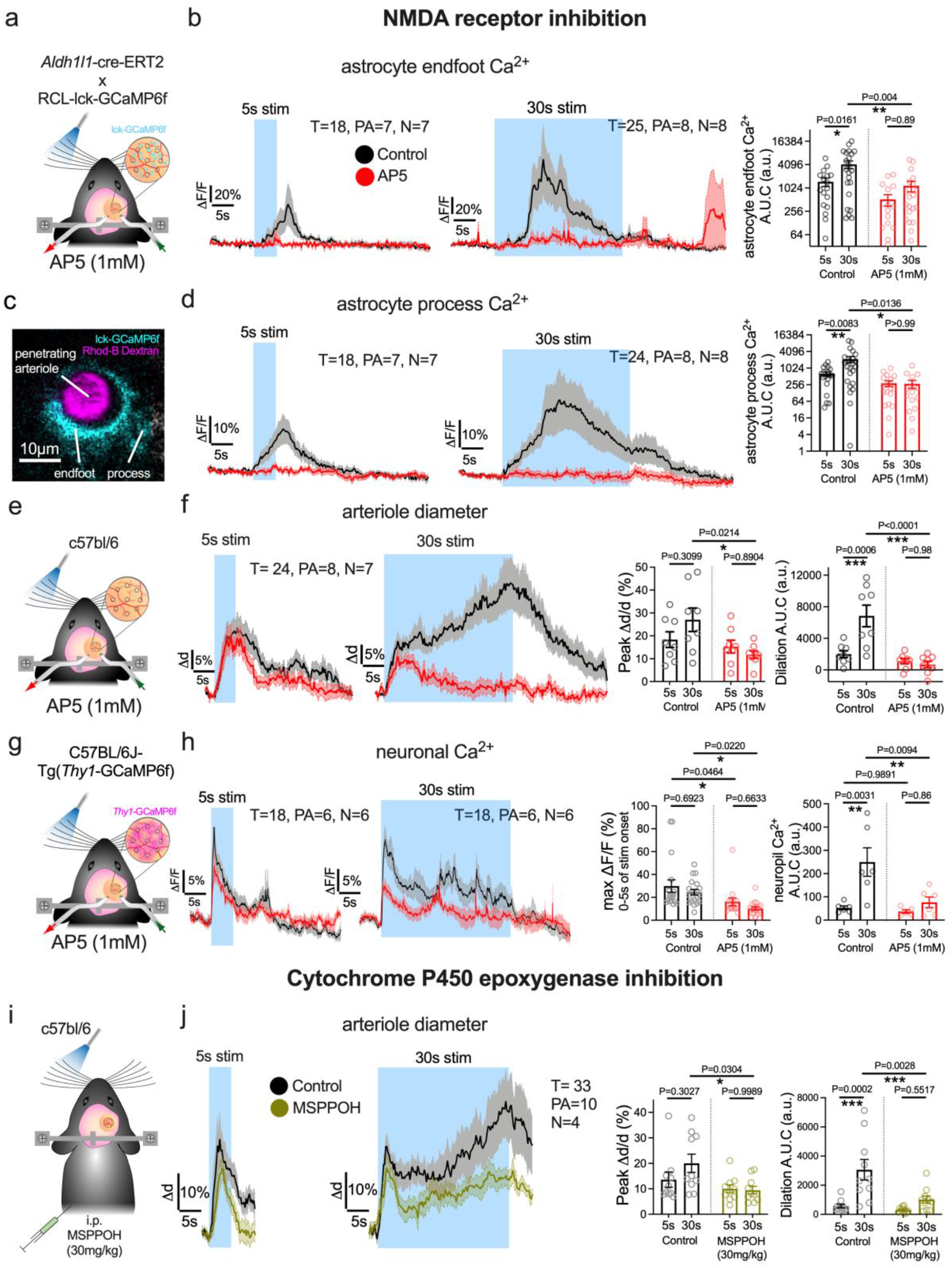
NMDA receptors and epoxyeicosatrienoic acids explain astrocyte-mediated amplification of sustained functional hyperemia. **a**) Cartoon of awake 2-photon imaging experiment with sensory stimulation and perforated window for AP5 superfusion in Aldh1l1-CreERT2 x RCL-lck-GCaMP6f mice. **b**) 2-photon image of a penetrating arteriole (red) and surrounding astrocyte expressing membrane targeted lck-GCaMP6f. **c**) *Left*: Average time series trace data of astrocyte endfoot Ca^2+^ in pre-drug control (black) and in the presence of AP5 (red) surrounding a penetrating arteriole in response to 5sec and 30sec whisker stimulation. *Right*: summary data of Area Under the Curve (AUC). **d**) Same as for *c* but for arbor astrocyte Ca^2+^. **e**) Cartoon of a C57Bl/6 mouse with whisker puffer and perforated cranial window for superfusion of AP5. **f**) *Left*: Average time series traces of penetrating arteriole diameter in pre drug control (black) or in the presence of AP5 (red), in response to 5sec or 30sec whisker stimulation. *Right*: summary data of dilation peak and area under the curve. **g**) Cartoon of a Thy1-GCaMP6f mouse with whisker puffer and perforated cranial window for superfusion of AP5. **h**) *Left*: Average time series traces of neuronal calcium in pre drug control (black) or in the presence of AP5 (red), in response to 5sec or 30sec whisker stimulation. *Right*: summary data of maximum neuronal Ca^2+^ signal in the first 5sec of stimulation and neuropil Ca^2+^ signal AUC of stimulation period. **i**) *Left*: Same as *e* but with sealed window and for i.p. delivery of the epoxygenase inhibitor MSPPOH. **j**) Left: same as *f* but showing pre-drug control (black) and in the presence of MSPPOH (green).

To test if NMDA inhibition abolished sustained neural activity as a potential explanation for the loss evoked astrocyte calcium elevation and late-phase functional hyperemia, we examined the temporal components of neuronal GCaMP6f during short and long duration whisker stimulation. We used C57BL/6J-Tg*(Thy1*-GCaMP6f) mice to measure neuronal Ca^2+^ signals before and during topical NMDAR antagonist D,L-AP5 (1mM) via a perforated coverglass window (Fig. 5g). We found that AP5 caused a reduction in the peak of the bulk neural calcium measurements around the arteriole to 5sec whisker stimulation (Initial Ca^2+^ peak: APV 16.1±3.4% vs control 29.8±5.4%, Two-way ANOVA and Tukey’s test: P=0.046, Fig. 5h) and also caused uniform reduction in the 30sec response (APV AUC:77±23 a.u. vs control AUC: 250±61 a.u., Two-way ANOVA and Tukey’s test: P=0.0094, N=5-6, Fig. 5h). While these data show AP5 effects sustained neural activity, the kinetic profile does not strongly affect the late phase over the early phase, suggesting other NMDAR-dependent neurovascular coupling mechanisms downstream of neural activity, such as astrocytes and perhaps endothelium ^60,62^ were affected.

Epoxyeicosatrienoic acids (EETs) have been demonstrated to contribute to functional hyperemia ^63–66^ and are primarily produced by astrocytes ^67,68^. Whether blocking EETs production has differential effects on brief vs sustained functional hyperemia has not been tested in awake mice. We measured functional hyperemia to 5sec and 30sec stimulation before and 1hr after i.p. injection of the epoxygenase inhibitor MSPPOH (30mg/kg) to c57bl/6 mice (Fig. 5i). Consistent with our proposed model, there was no difference in arteriole dilation to 5sec stimulation in the presence of MSPPOH (Δd/d=10±1.5%), compared to before (Δd/d=13.6±3%, Two-way ANOVA and Tukey’s test: P=0.7581) (Fig. 5j). However, 30sec stimulation was significantly attenuated by MSPPOH (Δd/d=9.5±1.5%) relative to control (Δd/d=20±3.6%, Tukey’s test: P=0.0304, PA=10, N=4 mice, Fig. 5j)(see Fig. 5j for AUC data). MSPPOH yielded a similar result examining short vs long afferent stimulation-induced arteriole dilation in acute cortical brain slices (Suppl. Fig. 9). These experiments show that EETs, likely downstream of NMDAR activation, are important to maintain sustained functional hyperemia.

## Discussion

By selectively attenuating or augmenting astrocyte Ca^2+^ signaling, we provide strong evidence in awake mice that astrocyte Ca^2+^ does not initiate or mediate functional hyperemia when neuronal activation is brief (<5sec) but is essential to amplify the CBF increase when neuronal activation is prolonged. Our work also provides new information on the origins of the bimodal neurovascular coupling response *in vivo*. Our data show that a non-neuronal cell type is late to ‘come online’ and at least partly explain the augmentation of the later phase. Interestingly, earlier work in awake animals showed that surface pial artery dilation to long stimulation was bimodal but the later phase was not amplified ^15,16^. As astrocytes do not directly influence surface pial arterioles, this is congruent with only observing amplification of sustained CBF increases at penetrating arterioles which are wrapped by astrocyte endfeet, and that this astrocyte-mediated response is not conducted upstream to the brain surface.

Previously, controversies about the role of astrocytes in functional hyperemia have arisen from challenges in achieving selective activation ^69^, lack of effect manipulating astrocyte IP3 signaling ^33–35,55,56^ and side effects of anesthesia/sedation ^13,70^. Increased neural activity is the main driver ^71^ and user of fresh O_2_^72^, yet the well-established linear correlation of BOLD signal to neural activity is lost with sustained brain activation ^73,74^. These results are consistent with a large body of literature showing delayed elevations in astrocyte Ca^2+^ when neural activity increases, as well as delayed increases in astrocyte metabolism ^75,76^. Our data show that the fully awake animal is necessary to observe sufficiency effects on arteriole diameter driven by astrocyte Gq ^33^, and suggest other IP3R subtypes play a role in CBF regulation. That selectively driving the astrocyte Gq pathway augmented only sustained functional hyperemia, which is impaired in a number of neurological conditions ^25–28^, suggests a potential therapeutic target to improve brain blood flow in these states.

Our *ex vivo* BAPTA and *in vivo* astrocyte membrane-tethered GCaMP6 transgenic mouse results, as well as CalEx data indicate that fast astrocyte Ca^2+^ signals that precede vasodilation ^19,21^ are not involved in the initiation or first dilation peak of functional hyperemia. These same data argue against fast acting astrocyte Ca^2+^ at the capillary, pre-capillary or sphincter level ^4,77,78^ driving a rapid conduction-mediated dilation to upstream arterioles ^79,80^. This is because both astrocyte CalEx *in vivo* and astrocyte BAPTA *ex vivo* reached these vascular compartments. While indirect effects of astrocyte Ca^2+^ clamp on neural activity could explain the decrease in sustained functional hyperemia ^81,41^, that 5sec evoked vasodilation was unaffected both *in vivo* and *ex vivo*, and that bulk neuronal Ca^2+^ measures directly around the arteriole *in vivo* were unchanged by CalEx, argues against this possibility. Furthermore, we found no impact of astrocyte Ca^2+^ clamp on evoked neural Ca^2+^ to long stimulation in brain slices, despite reduced vasodilation. Finally, a selective effect of the EETs synthesis blocker MSPPOH on the late phase, a putative NVC astrocyte pathway, further supports a direct role of endfeet to amplify late phase arteriole dilation. However, it was expected that astrocyte CalEx would have some impact on the local neural activity given the literature ^36,41^ and that CalEx targets to both perisynaptic processes as well as endfeet. Indeed, our detailed analysis of neuronal Ca^2+^ transients in the parenchyma showed some effects, yet without a preference for the late phase of 30sec stimulation. In either scenario (direct effect on the vessel or indirect effect via neural activity), our data points to an important role for astrocyte Ca^2+^ in this amplification phenomenon.

Whisker stimulation elevated locomotion and heightened arousal in our experimental paradigm. These behavioral states are often linked ^50^, and activating neuromodulatory pathways can facilitate astrocyte activation in the neocortex ^44,45^. Two of our results suggest that neuromodulatory systems contribute to functional hyperemia directly through astrocytes. First, the level of locomotion impacted only astrocyte endfoot Ca^2+^ transients and clamping these transients by CalEx significantly reduced late hyperemia ^82^. Second, selective activation of Gq signaling, a common pathway for neuromodulators in astrocytes, amplified late hyperemia without modifying neuronal Ca^2+^ responses. Identifying the link between behavior-related neuromodulatory inputs and astrocyte-derived vasoactive mediators is warranted.

Surprisingly, we found a temporally distinct contribution of NMDA receptors to eliciting delayed astrocyte Ca^2+^ transients and mediating the late phase of functional hyperemia. NMDA receptor activation likely controls astrocyte Ca^2+^ indirectly through diffusible mediators such as nitric oxide released from post-synaptic neurons, because astrocyte-patch based NMDA inhibition with intracellular MK-801 preserves evoked Ca^2+^ transients ^60^. Nevertheless, since the block of the late phase vasodilation by AP5 is much more pronounced than by astrocyte CalEx, and because AP5 caused a reduction in neuronal activity measured by GCaMP, part of the decrement in the late component of functional hyperemia may be attributed to decreased neuronal activity. It is interesting to speculate that the same synaptic NMDA receptors necessary for long-term potentiation and thus memory are the same for late phase hyperemia, which could have arisen due to the enhanced energy requirements of plasticity ^83,84^. However, late-phase hyperemia could also develop by NMDA receptor activation on endothelial cells. This process requires Ca^2+^-dependent D-serine release by astrocytes ^85,86^ which would be additionally inhibited by AP5 because this treatment abolished sensory-evoked astrocyte Ca^2+^ elevations.

In sum, these data 1) help clarify the literature on the role of astrocytes in functional hyperemia – a phenomenon that is fundamental to fueling the brain; 2) provide important insights into the cell-type specific underpinnings of bimodal neurovascular coupling in awake animals and 3) improve our understanding and interpretation of fMRI data which routinely employs stimulations or tasks of long duration.

## Materials and Methods

### Animals

All animal procedures were performed in accordance with guidelines approved by the Animal Care and Use committee of the University of Calgary (protocols AC19-0170 and AC19-0109). Experiments were performed on male P22-90 c57bl/6 mice (N=41: 5 acute window; 11 CalEx; 11 mutant CalEx control, 4 MSPPOH; 7 AP5; 3 thinned skull), transgenic *Aldh1l1*-Cre/ERT2 x RCL-GCaMP6s (Ai96)(N=9), *Aldh1l1*-Cre/ERT2 x R26-Lck-GCaMP6f (N=13), *PDGFRß*-Cre x RCL-GCaMP6s (N=11), *Aldh1l1*-Cre/ERT2 x CAG-LSL-Gq-DREADD (N=11), C57BL/6J-Tg*(Thy1*-GCaMP6f) (N=6). For the use of the Ai96 conditional line, three consecutive tamoxifen injections were administered (100 mg/kg, 10 mg/mL corn oil stock, Sigma, St. Louise MO USA) between the age of P19 to P45 and the awake imaging protocol followed at least 2 weeks later. The other ERT2 expressing mice received 4-hydroxy-tamoxifen for 2 consecutive days between P2-4 (16 mg/kg, 4mg/ml corn oil + 12 v/v (%) absolute ethanol) i.p. in a volume of 20μl.

### Two-photon imaging

Two custom-built two-photon laser-scanning microscopes were used, one of which was optimized for acute brain slices and patch-clamp electrophysiology ^87^ and another that was designed for awake *in vivo* experiments ^88^. Both microscopes were supplied by the same Ti:Sapph laser (Coherent Ultra II, 4 W average power, 670–1080nm, ∼80MHz) and equipped with the following parts: objectives (Zeiss 40x NA 1.0, Nikon 16x NA 0.8), a green bandpass emission filter (525–40nm) and an orange/red bandpass emission filter (605–70nm) coupled to photomultiplier tubes (GaAsP Hamamatsu). The open-source scanning microscope software ScanImage (version 3.81, HHMI/Janelia Farms) running in Matlab ^89^ was used for image acquisition. During acute brain slice experiments, we acquired time-series images at 0.98 Hz using bidirectional scanning of a 512 × 512 pixel-size area on a single focal plane visualizing fluorescent Ca^2+^ indicator-labeled cells and the center of a penetrating arteriole lumen in a depth of 40-120μm. For stimulation experiments, the Ti:Sapph was tuned to 850nm to excite Rhod-2/AM and FITC dextran fluorescent indicators. Filling of the astrocyte network with Alexa-488 was imaged at 780nm. *In vivo* experiments using GCaMP6 signals were imaged at 920-940nm, mCitrine at 980nm. Imaging astrocytes and vessels was performed at a standard 3.91Hz imaging frequency, neuronal Ca^2+^ in the CalEx and GqDREADD experiments at 7.82Hz and in the AP5 experiments at 15.64Hz. Fluorescent signals were recorded at 50-250μm depth of the sensory cortex.

### Acute brain slice preparation

We used Sprague-Dawley rats (p21-33) to prepare acute coronal slices of the sensory-motor cortex. Rats were anesthetized with isoflurane (5% induction, 2% maintenance). Two min after a fluorescent dye injection in the tail vein, animals were decapitated and their brains were quickly extracted and submerged into an ice-cold slicing solution (composition in mM: 119.9 N- methyl D-glucamine, 2.5 KCl, 25 NaHCO_3_, 1.0 CaCl_2_-2H_2_O, 6.9 MgCl_2_-6H_2_O, 1.4 NaH_2_PO_4_-H_2_O, and 20 glucose), continuously bubbled with carbogen (95% O_2_, 5% CO_2_) for 2 min. We cut 400μm thick coronal slices with a vibratome (Leica VT 1200S) and placed them to recover for 45min on a fine mesh in artificial cerebrospinal fluid continuously bubbled in carbogen at 34°C. ACSF contained the following (in mM): 126 NaCl, 2.5 KCl, 25 NaHCO_3_, 1.3 CaCl_2_, 1.2 MgCl_2_, 1.25 NaH_2_PO_4_, and 10 glucose.

The aCSF was bubbled with a gas mixture containing 30% O_2_, 5% CO_2_ and 65% N_2_ following the recovery for the dye loading period and during imaging experiments in order to enhance vascular dilation responses as described previously ^90^. We performed imaging at room temperature (22°C) and the imaging bath was perfused with aCSF at 2ml/min by a sealed, gas pressure driven system. The superfused aCSF was supplemented with the arteriole pre-constrictor U46619 (125nM) starting at least 30 min prior to each experiment to allow for the vessels to develop a steady tone. Drugs were infused into the main superfusion ACSF line (moving at 2mL/min) through a side line driven by a syringe pump at a rate of 0.2mL/min at ten-fold of the desired concentration to achieve the intended final concentration in the bath.

### Dye loading

We labelled penetrating arterioles with Fluorescein Isothiocyanate (FITC) dextran (2000 KDa, Sigma Aldrich) via a tail vein injection (15mg dissolved in 300μl lactated Ringer’s solution (5%)) under anesthesia immediately before slicing. Following slice recovery, slices were bulk loaded in Rhodamine-2 Acetoxymethyl Ester (Rhod-2/AM)(15μM, dissolved in 0.2% DMSO; 0.006% Pluronic Acid; 0.0002% Cremophore EL) in 3mL of aCSF while bubbled with a 30% O_2_-containing gas mixture (see above) in a small incubation well for 45min at 33°C. In acute slice experiments astrocytes were identified by a brighter baseline dye fluorescence than neurons due to their preferential Rhod-2/AM uptake and by the presence of endfeet on vasculature.

### Materials

FITC-dextran (2000KDa), rhodamine B isothiocyanate-dextran (Rhodamine-B-dextran, 70KDa), dexamethasone 21-phopshate disodium, Chremophore^®^ EL, DMSO were obtained from Sigma Aldrich. D- (-)-2-Amino-5-phosphonopentanoic acid (AP5) and Compound 21 dihydrochloride was purchased from Hello Bio Inc. (Princeton, NJ). U46619, N-(methylsulfonyl)-2-(2-propynyloxy)-benzenehexanamide (MSPPOH), medotomidine hydrochloride and atipamezole hydrochloride were ordered from Cayman Chemicals. BAPTA-tetrapotassium and Alexa 488 hydrazide were obtained from Invitrogen, isoflurane from Fresenius Kabi Canada Ltd., Pluronic F127 and Rhod-2/AM from Biotium, buprenorphine from Champion Alstoe Animal Health, and enrofloxacin from Bayer, meloxicam from Boehringer Ingelheim, ketamine from Vétoquinol N.-A. Inc. AAV2/5.GfaABC_1_D.GCamP6f and AAV9.hSyn.GCaMP6f were ordered from Penn Vector Core, PA, USA, pZac2.1-GfaABC1D-mCherry-hPMCA2w/b plasmid was ordered from Addgene.

### Electrical stimulation

The activation of afferent fibres was elicited by electric stimulation using an assembly of a Grass S88X stimulator, a voltage isolation unit and a concentric bipolar electrode (FHC). We softly positioned the electrode onto the surface of the slice by a micro-manipulator ∼250μm lateral to the penetrating arteriole. The tissue was stimulated for 5sec or 30sec with a theta burst pattern (trains of 1 msec mono-polar pulses at 100Hz for 50msec, repeated at 4Hz) using a 0.1V higher intensity than the threshold of astrocyte endfoot activation (1.1-1.3V). This threshold was determined by gradually ramping the voltage (+0.1V) of a low frequency (20Hz) 5sec stimulation until an endfoot Ca^2+^ transient was observed. For some experiments (see Suppl. Fig. 9), we employed a 20Hz stimulation for 5sec with low intensity that did not activate astrocyte endfeet and subsequent 30sec stimulations used the same voltage.

### Astrocyte patch clamping experiments

Patching experiments were conducted similar to our previous publication ^32^. Before the patching process, slices were electrically stimulated for 5 or 30sec with a theta burst pattern. Next, astrocytes were selected in the 30-70 μm proximity of an arteriole at a 30-60 μm depth, typically above the imaging plane of the arteriole (Suppl. Fig. 3a). A 3-5 MΩ resistance glass electrode formed a GΩ seal on the cell body membrane and when whole cell configuration was achieved astrocytes were filled with an internal solution containing the fluorescent dye Alexa-488 (100-200 μM) to visualize diffusion, and the following substances (in mM): 8 KCl, 8 Na-gluconate, 2 MgCl_2_, 10 HEPES, 4.38 CaCl_2_, 4 K-ATP, and 0.3 Na-GTP, 10 K-BAPTA (1,2-bis(o-aminophenoxy) ethane-N,N,N’,N’-tetraacetic acid) with 96 K-gluconate. Osmolality was adjusted to ∼285 mOsm, pH was corrected with KOH to 7.2. We voltage clamped astrocytes at -80 mV and confirmed the cell type by a low input resistance (10-20 MΩ) and dye filling of the endfeet circumventing a large section of the arteriole (Suppl. Fig. 3a-b). Importantly, dye filling of the endfeet with a clamping solution of 100nM Ca^2+^ did not change arteriole tone *per se* (see Suppl. Fig. 4). We allowed 15-30 min for any internal solution in the extracellular space from the patching process to washout from the tissue and for the arteriole tone to return to baseline. Then, while in stable whole-cell configuration, we repeated the electric stimulation to compare to the first response.

### Awake *in vivo* two-photon imaging of acute cranial window

Male mice were used according to the protocol previously developed by our laboratory ^88^. Briefly, on day 1, under isoflurane anesthesia (induction 4%, maintenance 1.5-2%), pain control (buprenorphine 0.05mg/kg) and antibiotic premedication (enrofloxacin 2.5 mg/kg), a light (0.5 g) titanium headbar was glued on the occipital bone under aseptic conditions with a three-component dental glue (C&B Metabond, Parkell Inc, NY, USA) and dental cement (OrthoJet Acrylic Resin, Land Dental MFG. CO., Inc., IL, USA). After a recovery period of 24hrs, the mice were trained on days 2 and 3 to habituate to imaging and air puff stimulation to the whiskers while head fixed but free to move on a passive, air supported spherical Styrofoam ball in a light tight environment (Fig. 1a). After 15 min of rest, 5sec continuous air puffs (30 psi, Picospritzer III, General Valve) were delivered to the contralateral whiskers 15 times every 10sec, 15sec, 30sec and 60sec, respectively on day 2 and 3. On day 4, a craniotomy was performed over the barrel cortex. Bone and dura were gently removed. For *Aldh1l1*-Cre/ERT2 x CAG-LSL-Gq-DREADD mice, Rhod-2 AM (15μM) in aCSF was incubated on the brain surface for 45min. Next, the window was fully sealed with a coverslip. In experiments where astrocyte endfeet were labelled with a red fluorescent dye, 0.2mL 5% FITC dextran solution was injected into the tail vein immediately before imaging, whereas Rhodamine-B dextran (0.2mL, 2.3%) was injected when astrocytes or neurons expressed GCaMP6. The animals were transferred to the imaging rig, where imaging started ∼30min after the animals awoke. Behavior was continuously monitored by a camera detecting the light of an infrared LED and was recorded during imaging. A 16x or a 40x water-immersion objective was positioned square to the surface of the window. Imaging was performed at 3.91Hz or at 0.49Hz when drug application was recorded. Rhod-2/AM only loaded astrocytes in mice *in vivo* (Fig. 4a) as neuronal compartments were neither labeled at rest nor appeared in response to stimulation ^88^.

### Awake *in vivo* two-photon imaging of chronic cranial window

Male, 3-4 months old c57bl/6 and *Aldh1l1*-Cre/ERT2 x R26-Lck-GCaMP6f mice were used. For head bar installation mice were premedicated with dexamethasone (2h prior, 6mg/kg s.c.). Under isoflurane anesthesia and aseptic conditions, the parietal, occipital and temporal bones were carefully separated from the surrounding soft tissue and a U-shaped metal headbar with 2 lateral arms was cemented to the occipital and temporal bones by a UV glue (Ivoclar vivident Tetric Evoflow). A conical cement wall was built around the right parietal cortex. The surgery was supported with antibiotic (enrofloxacin), analgesic (buprenorphen 1mg/kg s.c.) and anti-inflammatory (meloxicam 5mg/kg s.c.) medication. After a minimum of 5-day recovery, mice were premedicated with dexamethasone (2-4h prior), enrofloxacin, meloxicam, buprenorphine and anesthetized with a mixture of ketamine (100mg/kg) and medotomidine (0.5mg/kg) i.p. The mice were temperature controlled and inhaled 100% O_2_ during cranial window implantation. A 3mm circular area was removed from the parietal bone over the whisker barrel area, leaving the dura intact. To prevent bone regrowth, a custom-cut T-shaped coverslip (inner diameter: 3 mm; outer diameter: 3.5mm, central thickness: 250μm)(Laser Micromachining Ltd.) was sealed into the hole with UV cement. Anesthesia was reversed with atipamezole (0.5mg/kg s.c.). Post-operative supportive therapy included a heated cage with O_2_-enriched air (until the mouse turned active) and recovery jello for diet. For CalEx, and mutant CalEx control mice, a tiny slit was cut in the dura, through which the viral construct was injected into the barrel cortex before sealing the craniotomy. Six weeks later, mice were trained the same as in the acute cranial window experiments applying 5sec continuous air puffs to the whiskers. For ball movement and pupil diameter recording 2 near-infrared LEDs (780nm) and 2 infrared-sensitive cameras (Basler) were pointed at a small area of the Styrofoam ball and the right eye of the mouse, respectively. We recorded pupil + ball movement at 40Hz, while ball movement for 2P imaging was recorded at 62.5Hz with the software Pylon Viewer. An open-source pulse train generator (Sanworks, Pulse Pal v2) was configured to synchronize trigger signals between the cameras, LEDs, the Picospritzer III and the microscope.

### CalEx Plasmid construction

To generate the AAV astrocyte-targeted expression construct of HA-tagged CalEx, the DNA sequence encoding hPMCA2w/b was PCR amplified from pZac2.1-GfaABC1D-mCherry-hPMCA2w/b (a gift from Baljit Khakh (Addgene plasmid # 111568 ; http://n2t.net/addgene:111568 ; RRID:Addgene_111568) with primers designed to incorporate a N-terminal HA tag encoding the amino acids YPYDVPDYA and used to replace mCherry-PMCA2w/b in the same backbone at the 5’ NheI and 3’ XbaI restriction sites using the NEBuilder Hifi DNA assembly kit, generating pZac2.1-GfaABC1D-HA-hPMCA2w/b ^23^. For the control construct, the critical Ca^2+^ binding site residue glutamic acid 457 was converted to Alanine using PCR site-directed mutagenesis, generating a mutant control CalEx (hPMCA2w/b (E457A)) incapable of binding or membrane transport of Ca^2+ 37,38^. The DNA sequences of all constructs were verified by Sanger DNA sequencing.

### AAV production

AAV viral vectors were generated using the methods of Challis et. al. ^91^. Briefly, 293FT cells (Thermofisher) were grown to ∼90% confluency in Corning hyperflasks (Corning) and co-transfected with 129 μg pHELPER (Agilent), 238 μg rep-cap plasmid encoding AAV5 capsid proteins (pAAV2/5 was a gift from Melina Fan (Addgene plasmid # 104964 ; http://n2t.net/addgene:104964 ; RRID:Addgene_104964) and 64.6 μg of either the CalEx plasmid (pZac2.1-GfaABC1D-HA-hPMCA2w/b) or control (pZac2.1-GfaABC1D-HA-hPMCA2w/b (E457A)) using the PEIpro transfection reagent (Polyplus). The AAV viral vector purification, concentration and titer determination was performed as previously described ^91–93^.

### AAV CalEx injection

A viral cocktail containing AAV2/5.GfaABC_1_D GCaMP6f (3 × 10^12^ gc/ml, Addgene) for astrocyte Ca^2+^ detection or AAV9.hSyn.GCaMP6f (3 × 10^12^ gc/ml, Addgene) for neuronal Ca^2+^ detection was mixed with the astrocyte-targeted CalEx encoding construct AAV2/5.GfaABC1D-HA-hPMCA2w/b (1.18 × 10^13^ gc/ml) in a 1:3 ratio and delivered by a Nanoject III (Drummond Scientific, Broomall, PA, USA) in a volume of 400nL. For the best possible control, we designed a mutated version of the CalEx virus in which changing the critical glutamic acid residue in TM4 (E457) to Alanine to render the pump incapable of transporting Ca^2+^. Training and imaging took place 6 weeks later, because this is sufficient time to achieve functionally effective CalEx expression ^36^.

### Immunohistochemistry

Slices were collected from control and CalEx mice at the end of the *in vivo* experiments and post-fixed in 4% paraformaldehyde overnight and then cryoprotected in 30% sucrose for another 24hr. Sections were cut on a cryostat to 30μm and free floated in PBS. The sections were first washed in PBS (phosphate buffered saline) for 20min, followed by 3 × 5 min washes in PBS; agitated at room temperature. Sections were permeabilized in 0.1% Triton X in PBS for 10 minutes. Anti-HA (rabbit) primary antibody (1:100: Abcam) was diluted in 2% BSA in PBS and incubated for 1 hr and agitated at room temperature. After primary antibody incubation, sections were washed three times with PBS and then incubated for 1hr with Alexa Fluor 555 (1:1000: Invitrogen) diluted in 2% BSA in PBS and agitated at room temperature. Sections were then washed again three times with PBS before mounting coverslips with ImmunoMount (Epredia) and read on a confocal fluorescence microscope (Leica TCS SP8, Leica Microsystems).

### Functional hyperemia

Two-photon imaging session started by confirming Rhod-2/AM, GCaMP6 or Gq DREADD-linked mCitrine expression and identifying the vascular network by z-stack imaging to a depth of ∼300μm with a 16x objective. Pial arterioles were distinguished from veins by spontaneous vasomotion and by positive responses to contralateral whisker air puff. Penetrating arteriole branches of the pial arteries were tracked and imaged at a depth of 50-250μm with a 40x or a 16x objective. Air puff was delivered through 2 glass capillary tubes positioned to deflect as many whiskers as possible without blowing air on the face (Fig. 1a). A 10sec baseline period was recorded after which a continuous or an intermittent (4Hz, 125msec pulses) 5sec or 30sec air puff (30psi) was employed. Stimulation was repeated 3-6 times and averaged for each arteriole. In case of chronic cranial window experiments, on the day before two-photon imaging we mapped the cortical center of activation to whisker stimulation by collecting intrinsic optical signals of hemoglobin under the cranial window. We flashed green light with a stable light-emitting diode (M530L3, Thorlabs Inc.) equipped with filtering (Semrock) and collimation optics (Thorlabs) at 10Hz and collected reflected light with a Basler Ace U acA1920-40 μm camera with Fujinon HF50XA-5M lens(pixel size = 5.86μm), during 30sec whisker stimulations. Penetrating arterioles for two-photon imaging were selected from a region where green light reflectance showed a major decrease during the period of whisker stimulation overlapping with the expression of the viral construct (good neuronal or astrocytic GCaMP6 expression). We measured pupil diameter changes in a separate set of trials with no stimulation, 30sec ipsilateral air puff to the body, or 30sec whisker stimulation while the pupils were constricted to mid-size with a low intensity continuous green light in the background.

### Data processing and statistical analysis

Ca^2+^ signals were analyzed in ImageJ (NIH) and calculated in Prism software (GraphPad Inc., La Jolla, CA) as follows: ΔF/F=((F_1_-F_0_)/F_0_) x 100, where F is fluorescence, 1 is at any given time point, and 0 is an average baseline value of 60sec *ex vivo* or 2sec *in vivo* pre stimulus. Ca^2+^ responses in brain slices were obtained by selecting a region of interest (ROI) and running the ‘intensity vs. time monitor’ plugin over the following compartments: (1) neuropil was outlined as an acellular region next to the region of vasodilation on the side closer to the stimulating electrode, (2) neuron soma responses were the average of 5 neuronal cell bodies visible in the imaging plane closest to the arteriole, (3) astrocyte soma signal was the average of 1-3 cells closest to the arteriole, (4) astrocyte endfoot was chosen as a clearly visible continuous structure adjacent to the area of vasodilation. The same ROIs were used after patching *in vitro* or after drug administration *in vivo*. Astrocyte GCaMP6 signals of endfeet and processes in all experiments were outlined by including regions around the arteriole where an increase in the fluorescent signal was visible after stimulation onset. On neuronal GCaMP6 recordings ROI was determined as a 5000μm^2^ polygonal perivascular area avoiding the shadow of the proximal section of the vessel.

In the absence of stimulus-evoked fluorescence changes, areas of basal GCaMP6 fluorescence were selected. Ca^2+^ signals are labeled as ΔF/F_c_ in control conditions, ΔF/F_BAPTA_ after BAPTA loading. Pre-processing of intraluminal dye recordings in ex vivo slices included xy-motion correction, median filter (0.5-2 pixel radius) and Gaussian filter (1-4 pixel radius) performed by ImageJ plugins. Intraluminal area was selected by intensity thresholding and quantified using the ‘Analyze particles’ function of ImageJ.

To determine the cross-sectional area of intraluminal dye-loaded arterioles *in vivo*, we used 3 independent methods to outline the edges of the vascular lumen. For Gq-DREADD (Fig. 4e-f) and MSPPOH experiments (Fig. 5i) we used manual outlining of individual images via Amazon Mechanical Turk (see section below), for thinned skull, and acute window experiments (Fig 1a-h, Fig. 5a-g, Suppl. Fig.1a), we used ImageJ analysis (see above). For CalEx experiments and Aldh1l1-Cre/ERT2 x R26-Lck-GCaMP6f chronic window experiments (Suppl. Fig 2d-f, Fig. 2) we averaged the results of the ImageJ analysis and of a custom-made automated luminal area tracking MATLAB script (described in detail in Haidey et al., 2021 Cell Rep ^94^) for each trial. Briefly, based on user defined grayscale input templates of luminal edges (ROIs), an area mask is computed by *inpolygon* in MATLAB to yield a binary matrix for each ROI template. The matrix of the input templates is used by Thirion’s DEMONS algorithm to find a distortion function that creates a cross-sectional area from the binary matrix for each frame. A colour-coded image of the distorted input template with the detected edges is shown in Suppl. Fig. 10i. Arteriole diameter changes were quantified as the relative change in the area of intraluminal dye along the longitudinal section (in slice) or calculated from the cross-sectional area (*in vivo*) of the vascular lumen, assuming a circular shape, as follows: diameter = 2x √(area/π). Stimulation-evoked vasodilation was first calculated as Δ diameter (Δd) = (diameter_1_-diameter_0_)/diameter_0_) x 100 (%), where diameter_1_ is the diameter of the vascular lumen at any time point and diameter_0_ is the average baseline vascular lumen diameter 2sec prior to stimulation onset. Response latency was determined as the time from stimulation onset until 2 consecutive values exceeded 3 standard deviations of the 2sec baseline. All peak responses during topical C21 application (first 10min from Ca^2+^ increase onset) were compared to the peak value of the 500sec baseline period (see Fig. 4c). AUC values were calculated from Δd traces as the summary of all positive peaks over the baseline value for 35min from stimulation onset.

Event-based analyses of Ca^2+^ signals in neuronal compartments were performed by an automated analysis toolkit (https://github.com/yu-lab-vt/AQuA) ^41^ on 5000μm^2^ peri-arteriole ROIs of image stacks recorded at 7.82Hz. A Gaussian spatial filter was applied blindly to each trial (1-4 standard deviation of the mean) to optimize the fidelity of event detection. Movement-related event detection artefacts were excluded. Absolute event frequency (calculated as the average number of events for each sec), peak fluorescence value (dFF Max), integrated Ca^2+^ event value (AUC), event area size (μm^2^) and the duration (time of rise and fall between the 10% of peak) were summarized from the analysis output.

Locomotion analysis: Recordings of a small area of the Styrofoam ball were analyzed by a custom MATLAB script to quantify ball movement on a relative locomotion scale where ‘0’ is assigned to the least movement (stationary ball) and 1 is the highest speed an animal reached during the experiment.

To quantify locomotion activity from recordings of a small region of the Styrofoam ball, we implemented a simple algorithm in ImageJ: briefly, the image stack was read in and duplicated; the first frame from ‘Stack1’ and the last frame from ‘Stack2’ were deleted; a pixel-by-pixel subtraction was performed for every image in the stack (i.e. Stack1 – Stack2); mean gray value was plotted against time for the resulting stack.

Pupil diameter analysis: Pupil diameter changes were recorded at 40Hz using an infrared camera mounted close to the animal’s eye. To quantify pupil diameter from the videos, we used single-animal DeepLabCut to train a model to track four points on the circumference of the pupil (two along the vertical axis and two along the horizontal axis). The following parameters were used: TrainingFraction=0.95; net_type=resnet_50; augmenter_type=imgaug; max_iters=500000. Tracking performance was assessed by visual examination of the processed videos and the filtered position traces; frames where the animal blinked (<0.5%) were discarded. The average of the horizontal and vertical distance was used as pupil diameter. A custom MATLAB script was used to determine pairwise distances and perform median filtering of the diameter traces.

Statistical analyses were performed on averaged trials of PA responses and on Ca^2+^ responses of individual stimulation trials *in vivo*. Normality was tested with Shapiro-Wilk test. Normally distributed data were evaluated with a paired or unpaired Student’s t-test, and non-normally distributed with Mann-Whitney or Wilcoxon test. The non-parametric Friedman test and Dunn’s multiple comparison test was applied to compare *in vivo* peak diameter changes on stimulation length (1- 5- 30sec). Response latency of arteriole, endfoot and process Ca^2+^ was analyzed with Kruskal-Wallis test and Dunn’s multiple comparison test (Suppl. Fig. 2d). Repeated-measures two-way ANOVA was run to compare the effects of stimulation length (5sec vs 30sec as within subject variables) and external manipulations (CalEx, or drug treatments), followed by Tukey’s post hoc test for pairwise comparison. Baseline arteriole diameter (intravascular area) changes were analyzed with the two-tailed non-parametric Wilcoxon test in Prism. Data are expressed as mean ± standard error of the mean (SEM). We analyzed the effect of locomotion as a co-variate for each trial on group differences in arteriole diameter, astrocyte and neuronal Ca^2+^ signals of the CalEx experiment using generalized linear models in SPSS (IBM SPSS Statistics, Version 25).

### Amazon Mechanical Turk analysis

Arteriole diameter changes of the CalEx experiments were analyzed by manual outlining of the FITC dextran loaded vascular lumen frame by frame in an outsourced and randomized fashion, using Amazon Mechanical Turk (Suppl. Fig. 10). It is an online crowdsourcing website, through which remotely located “crowd-workers” are hired to perform a discrete on-demand task. A custom-made GUI (https://github.com/leomol/vessel-annotation) facilitated the manual outlining of the vascular cross-section over the course of the recording (https://codepen.io/leonardomt/full/jOEvPvY). An annotation task consisted of adjusting a polygonal shape with a fixed number of vertices on the top of the vessel. To keep annotations consistent, crowd-workers could inspect the time evolution of the vessel and their annotations by freely scrolling through the time-lapse images. Annotation tasks were randomly assigned to 10 different crowd-workers, blind to the experiment. Each stack of images was then visually examined and accepted or rejected based on whether the instructions were followed or not, i.e., if outlines were not consistently labeled over time or if there was a clear mismatch in the outlines with respect to the brain vessel. The area of each polygonal shape was then calculated and normalized (Suppl. Fig. 10).

## Acknowledgement

This work was supported by the Canadian Institutes of Health Research. G.R.G. was supported by Canada Research Chairs. We appreciate Dr. Marine Tournissac’s assistance in establishing the chronic cranial window model. We thank Dr. David Rosenegger for preliminary patch experiments. We thank Dr. Ciaran Murphy-Royal, Brian MacVicar and Rebecca Williams for their valuable scientific comments. We thank the CSM Optogenetics Facility for hosting the web services for crowdsourced annotations. We are grateful for Dr. Frank Visser for transgenic mouse sample genotyping. We thank the developers and distributors of ScanImage open-source control and acquisition software for two-photon laser-scanning microscopy.

## Author’s contribution

A.I. designed and performed experiments, analyzed results and wrote the paper. M.V. and G.P. performed experiments and analyzed results, C.C. analyzed results, C.H.T. performed experiments, X.Y., and B.S.K., designed CalEx virus construct, F.V. designed the mutant CalEx virus construct, C.B. did immunohistochemistry, L.M. designed platform for crowdsourcing annotations with Amazon Mechanical Turk and supervised analysis. G.R.G. designed experiments and wrote the paper.

