## Supplemental figures and legends for "Astrocytes amplify neurovascular coupling to sustained activation of neocortex in awake mice"

### Supplemental figures and legends, Institoris A et al.

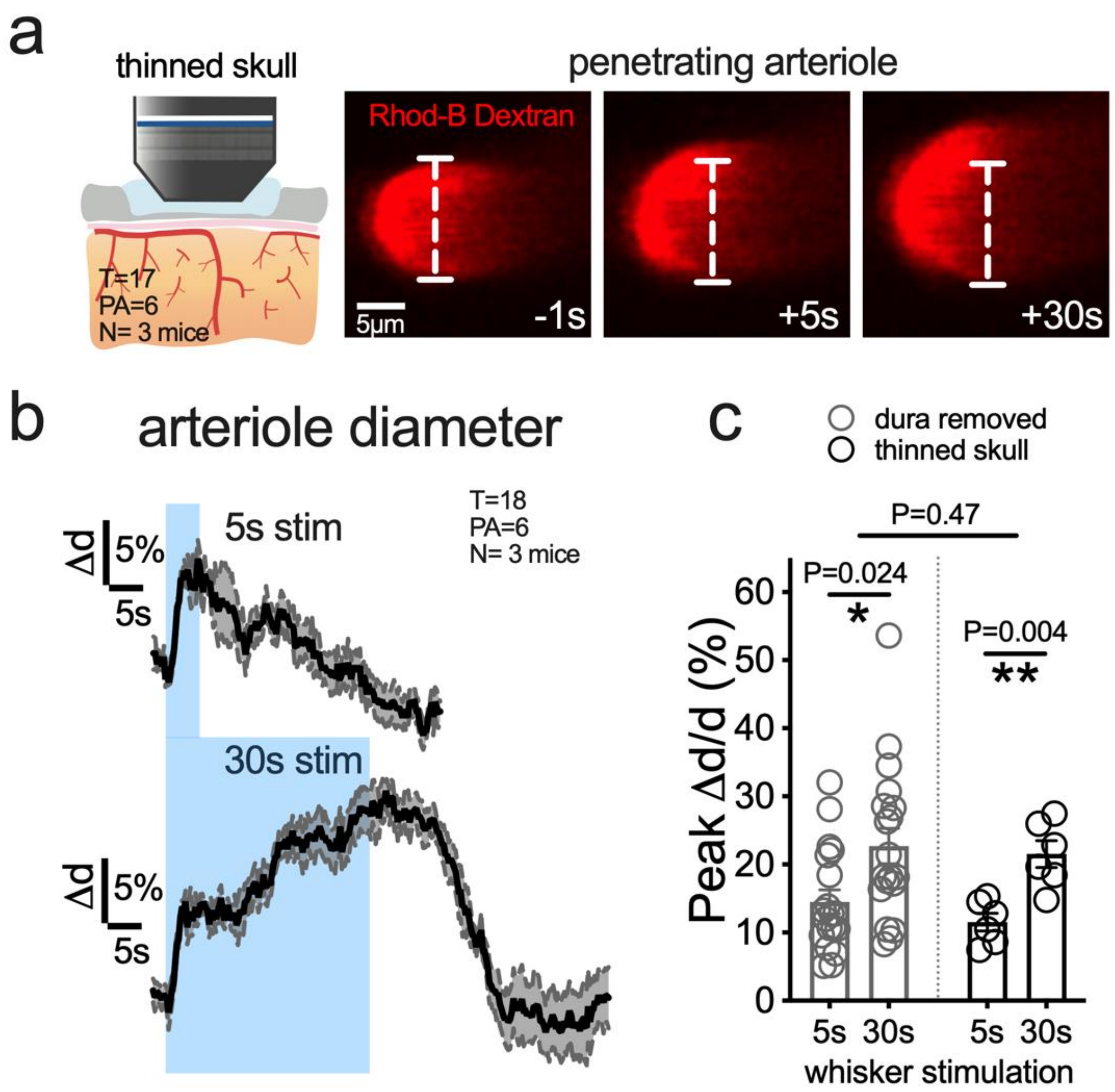


**<u>Supplemental Figure 1</u>: The bimodal functional hyperemia response is preserved after dura removal. a**) Cartoon of the experimental preparation (left) and 2P images of i.v. Rhodamine-B-dextran labelled penetrating arteriole before and during 30sec whisker stimulation imaged through an acute thinned skull preparation. **b**) Averaged traces of arteriole dilation to 5sec (upper) and 30sec (lower) whisker stimulation shows a bimodal arteriole response to 30s stimulation in a thinned skull preparation. **c**) Summary data of peak diameter changes of penetrating arterioles (trials averaged) comparing responses of the thinned skull prep to the responses of dura removed preparation.

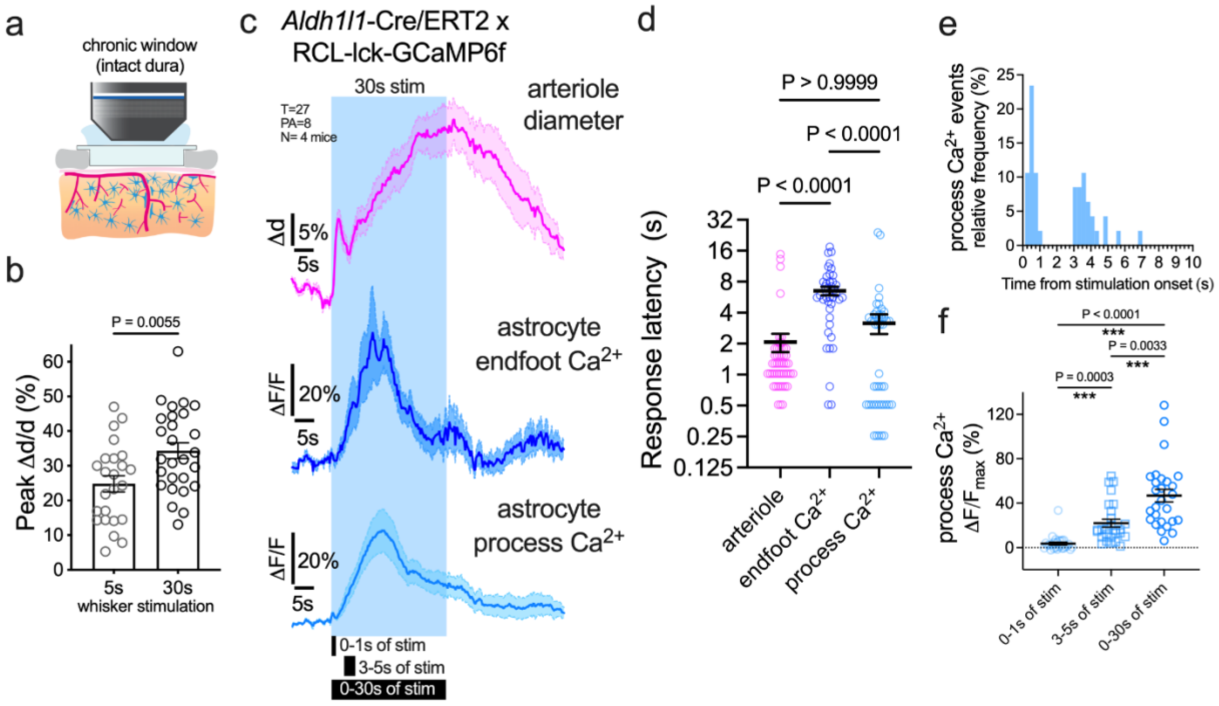


**<u>Supplemental Figure 2</u>: Astrocyte $Ca^{2+}$ signals of membrane tethered GCaMP6f during sustained functional hyperemia. a**) Cartoon of the chronic cranial window preparation implanted with a T-shaped circular coverslip over the dura. **b**) Averaged traces of arteriole dilation to 5sec and 30sec whisker stimulation (unpaired t test; T=24-27, PA=8-9, N=4). **c**) Arteriole diameter (top), astrocyte endfoot $Ca^{2+}$ (middle) and fine process $Ca^{2+}$ (bottom) responses to 30sec whisker stimulation. **d**) Summary of response onset (calculated from each trial as 3 x SD above baseline) for dilation and astrocyte $Ca^{2+}$ (Kruskal-Wallis test: P<0.0001 with Dunn's post hoc comparison). **e**) Relative frequency histogram of process $Ca^{2+}$ events from stimulation onset reveals an ultrafast (0-1sec) and a delayed (3-5sec) population. **f**) The size of astrocyte process $Ca^{2+}$ rise for ultrafast, delayed and overall (0-30sec) signals during stimulation (Friedman test: P<0.0001 with Dunn's post hoc comparison). All data are mean±SEM.

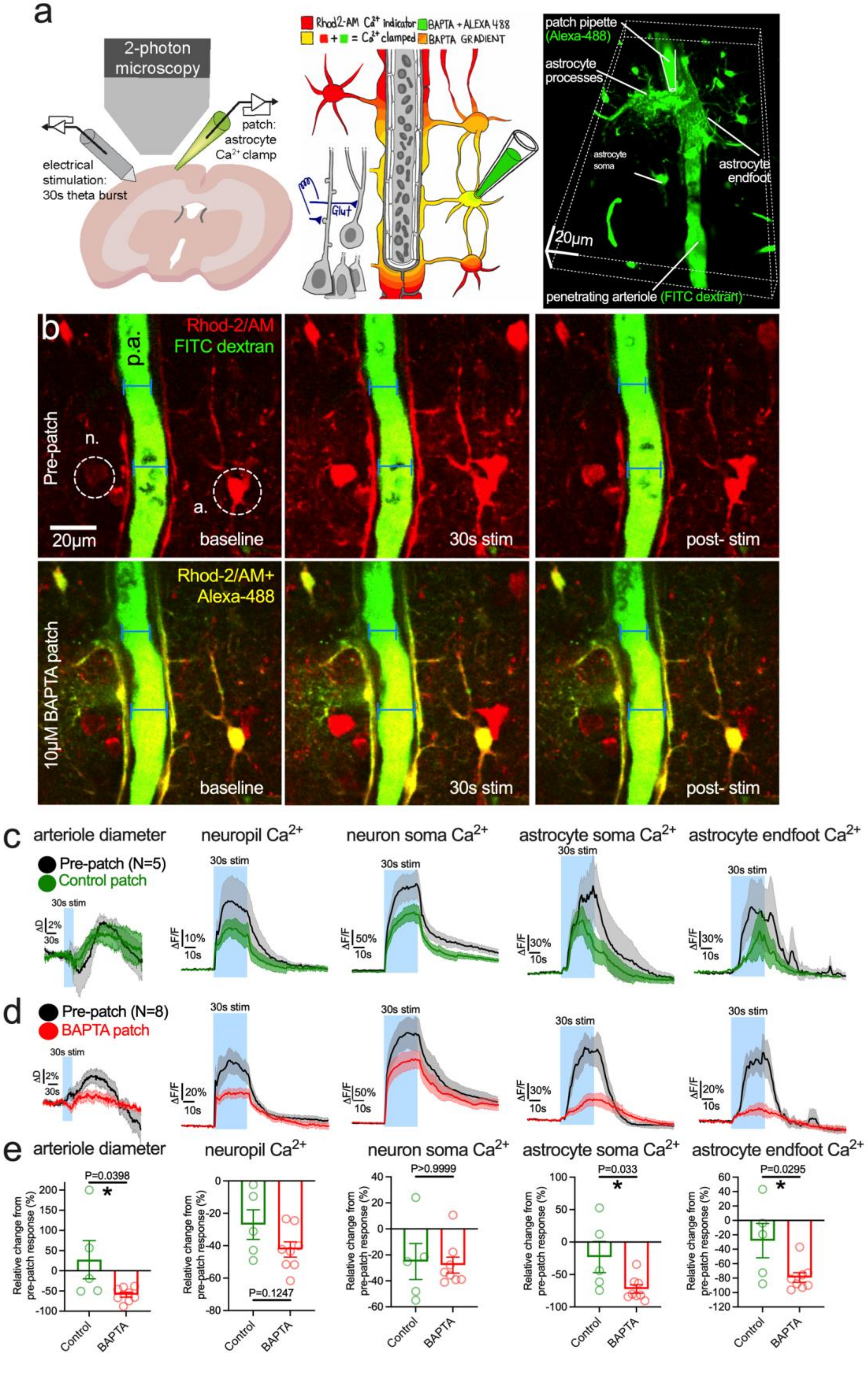
a
2-photon microscopy
electrical stimulation: 30s theta burst
patch: astrocyte Ca2+ clamp
Rhod2-AM Ca2+ indicator
BAPTA + ALEXA 488
= Ca2+ clamped
BAPTA GRADIENT
Glut
patch pipette (Alexa-488)
astrocyte processes
astrocyte soma
astrocyte endfoot
20μm
penetrating arteriole (FITC dextran)
b
Rhod-2/AM
FITC dextran
p.a.
Pre-patch
n.
a.
20μm
baseline
30s stim
post- stim
Rhod-2/AM+ Alexa-488
10μM BAPTA patch
c
arteriole diameter
neuropil Ca2+
neuron soma Ca2+
astrocyte soma Ca2+
astrocyte endfoot Ca2+
Pre-patch (N=5)
Control patch
30s stim
d
Pre-patch (N=8)
BAPTA patch
e
P=0.0398
P=0.1247
P>0.9999
P=0.033
P=0.0295
Relative change from pre-patch response (%)
Control
BAPTA

**Supplemental Figure 3: Astrocyte $Ca^{2+}$ clamp in brain slices with patched BAPTA reduces arteriole dilation to 30sec of high frequency afferent stimulation**. **a**) cartoon of experimental brain slice setup with patch infusion of BAPTA into the astrocyte network. Image on the right is a rotated z-stack of astrocytes patch-filled with Alexa-488 hydrazide around a FITC-dextran labelled arteriole. **b**) Upper: Image time series showing astrocyte $Ca^{2+}$ elevation and dilation to 30sec of theta burst electrical stimulation of afferents. Lower: the same stimulation is given in the presence of astrocyte network $Ca^{2+}$ clamp (yellow astrocytes) and vasodilation is blocked. **c** and **d**) Average time series traces in response to 30 sec of afferent stimulation showing arteriole diameter, neuropil $Ca^{2+}$, Neuron soma $Ca^{2+}$, astrocyte soma $Ca^{2+}$ and endfoot $Ca^{2+}$. Control, pre-patch traces are shown, followed by a patch infusion of a control internal solution (upper) or a $Ca^{2+}$ clamp internal solution containing BAPTA (lower). **e**) Summary data of percent changes from the pre-patch responses to either the control patch or the BAPTA patch condition. These data show that only the reduction in astrocyte $Ca^{2+}$ can explain the loss of dilation to 30sec stimulation in the astrocyte BAPTA patch condition.

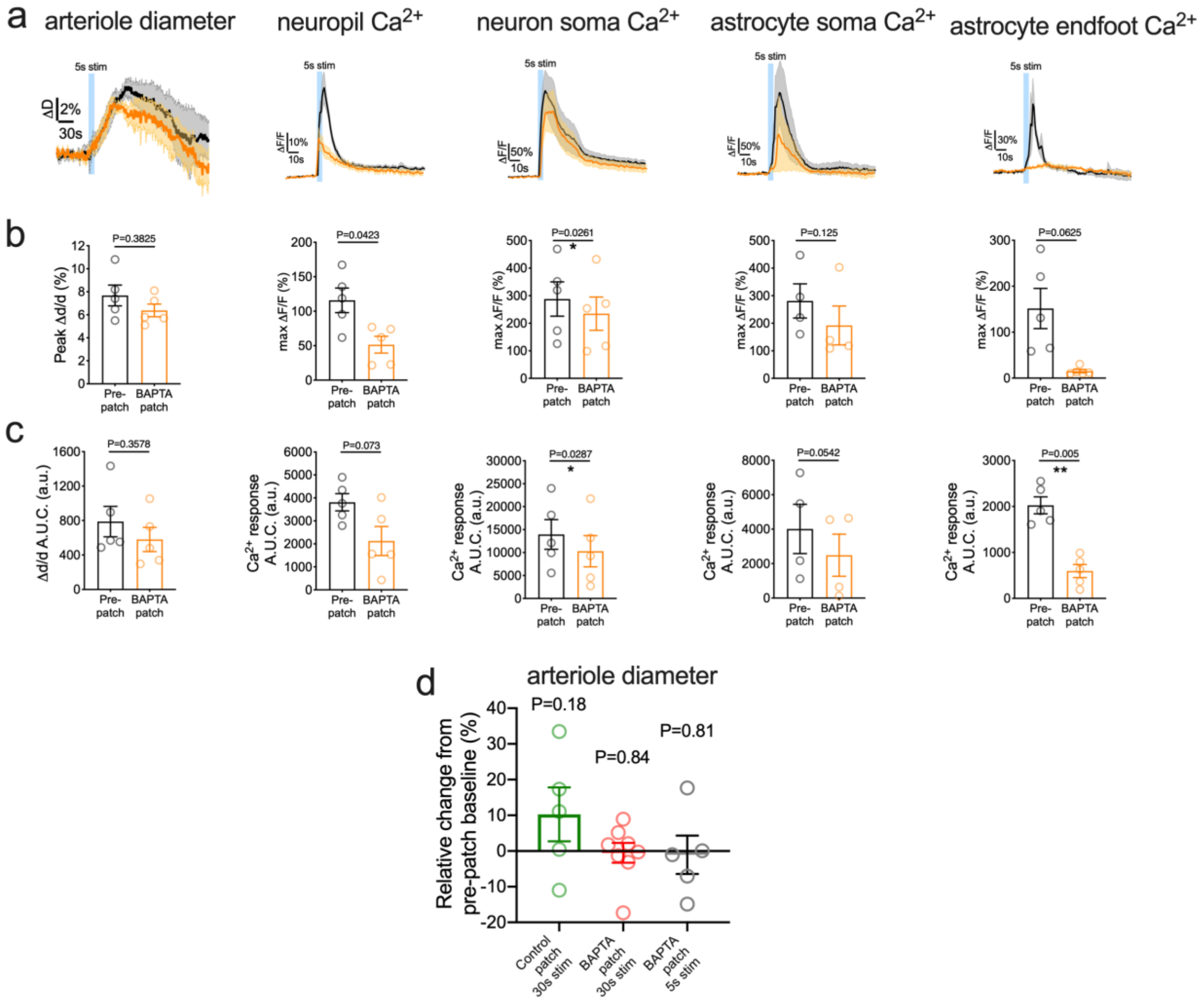


<u>Supplemental Figure 4:</u> **Astrocyte $Ca^{2+}$ clamp in brain slices with patched BAPTA has no effect on evoked arteriole dilation to 5sec high frequency afferent stimulation**. **a**) Average time series traces in response to 5 sec of theta burst afferent stimulation in control astrocyte patch (black) or $Ca^{2+}$ clamp patch (orange) showing arteriole diameter, neuropil $Ca^{2+}$, Neuron soma Ca2+, astrocyte $Ca^{2+}$ and endfoot $Ca^{2+}$. **b** and **c**) Summary data of peak response, or area under the curve (AUC) is shown underneath each set of traces. Paired t-test or Wilcoxon test was used. **d**) Summary data showing that neither the control patch internal solution, nor the $Ca^{2+}$ clamp internal solution to 100nM free $Ca^{2+}$ (in the 5sec and 30sec experiments) affected resting arteriole diameter after the 15min whole-cell equilibration period. Paired t-test between pre- and post-patch arteriole baseline diameter.

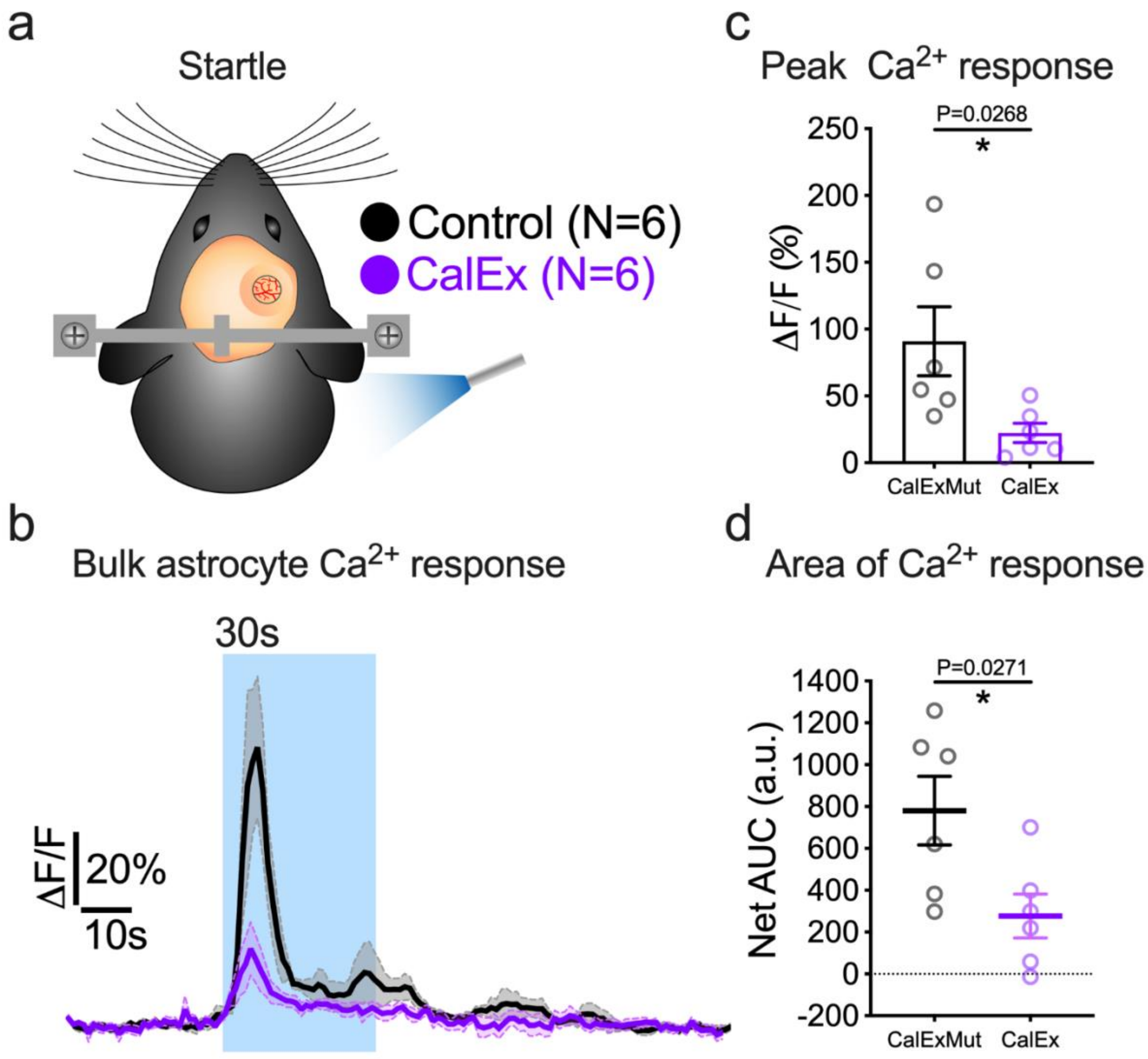


<u>Supplemental Figure 5</u>**: Expression of astrocytic plasma membrane $Ca^{2+}$ ATPase (CalEx) decreases the evoked $Ca^{2+}$ response to startle**. **a**) Cartoon of experimental setup using an untrained body air puff to startle the mouse. **b**) Average time series curves of astrocyte $Ca^{2+}$ in response to startle, with CalEx and GCaMP6f AAV vs control AAVs. **c**) Summary data of peak $Ca^{2+}$ response. **d**) Summary data of area under the curve (AUC) $Ca^{2+}$ response.

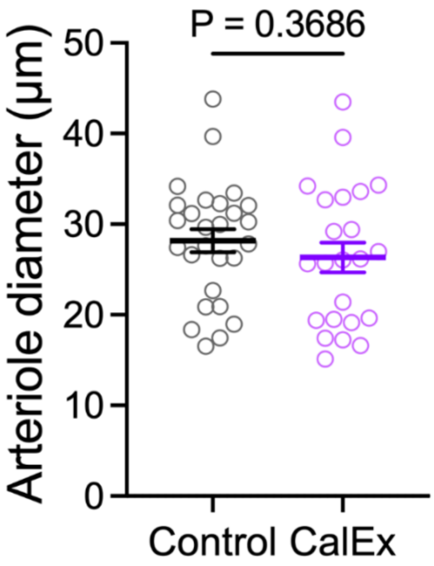


**<u>Supplemental Figure 6</u>: CalEx did not change baseline penetrating arteriole (PA) diameter.** Baseline arteriole diameter of Control (PA=27, N=11) and CalEx (PA=23, N=10) arterioles calculated from averaged 10sec pre-stimulus baseline recording (3-7 trials per PA).

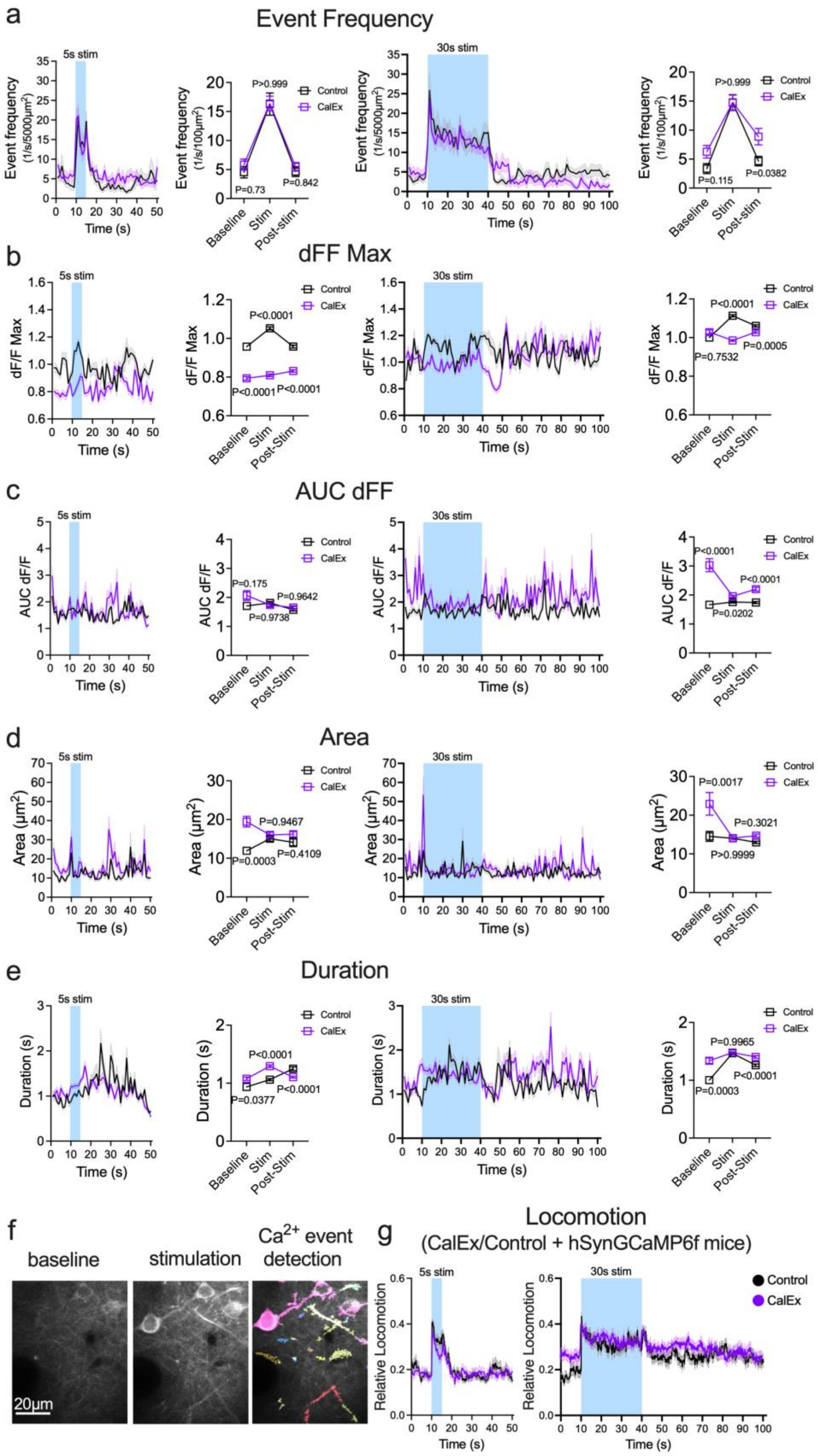
a
Event Frequency
5s stim
Event frequency (1/s/5000µm²)
Time (s)
Event frequency (1/s/100µm²)
P>0.999
P=0.73
P=0.842
Control
CalEx
Baseline
Stim
Post-stim
30s stim
P=0.115
P=0.0382
b
dFF Max
dF/F Max
P<0.0001
P=0.7532
P=0.0005
Post-Stim
c
AUC dFF
AUC dF/F
P=0.175
P=0.9642
P=0.9738
P=0.0202
d
Area
Area (µm²)
P=0.9467
P=0.0003
P=0.4109
P=0.0017
P=0.3021
P>0.9999
e
Duration
Duration (s)
P=0.0377
P=0.9965
f
baseline
stimulation
Ca²⁺ event detection
20µm
g
Locomotion
(CalEx/Control + hSynGCaMP6f mice)
Relative Locomotion

**<u>Supplemental Figure 7</u>: Automated $Ca^{2+}$ event detection analysis shows neuronal $Ca^{2+}$ differences are unrelated to CalEx effect on arteriole. a**) Left: Absolute neuronal (soma + neuropil) $Ca^{2+}$ event frequency curves (1sec binning of 7.81Hz recording) and averaged event frequencies of baseline, stimulation, and post-stimulation periods for 5sec whisker stimulation. Right: Same but for 30sec stimulation. Same layout for panels b-e. **b**) Summary curves and averaged values of baseline, 5sec (Left) and 30sec (Right) stimulation and post-stimulation periods for maximal relative fluorescence of individual events (Max dF/F) show significantly larger $Ca^{2+}$ peaks for control than for CalEx-injected mice (Two-way ANOVA with Tukey's test) but not during the later phase of 30sec stimulation. **c**) Area Under the Curve (AUC) of individual neuronal $Ca^{2+}$ event-related fluorescence changes (dF/F) demonstrate larger signals during baseline in the CalEx group. **d**) Area (size) of individual neuronal $Ca^{2+}$ events are also larger at baseline for CalEx than control. **e**) The average duration of $Ca^{2+}$ events for CalEx-injected mice are overall longer than for control virus injected mice except during sustained stimulation. **f**) Raw 2-photon image of neuronal $Ca^{2+}$ events in GCaMP6f expressing neuronal structures in layer 2 of the barrel cortex (Left) before and (Middle) during whisker stimulation. Right: Colour-coded detection of individual $Ca^{2+}$ events by an automated $Ca^{2+}$ event detection toolkit (https://github.com/yu-lab-vt/AQuA). **g**) Summary of relative locomotion curves for 5sec (Left) and 30sec (Right) whisker stimulation in mice injected with CalEx or its mutant control virus mixed with an AAV-hSynGCaMP6f virus indicate similar locomotion pattern during 30sec stimulation. Locomotion differences at baseline and 5sec stimulation between control and CalEx could account for the differences in individual $Ca^{2+}$ event properties.

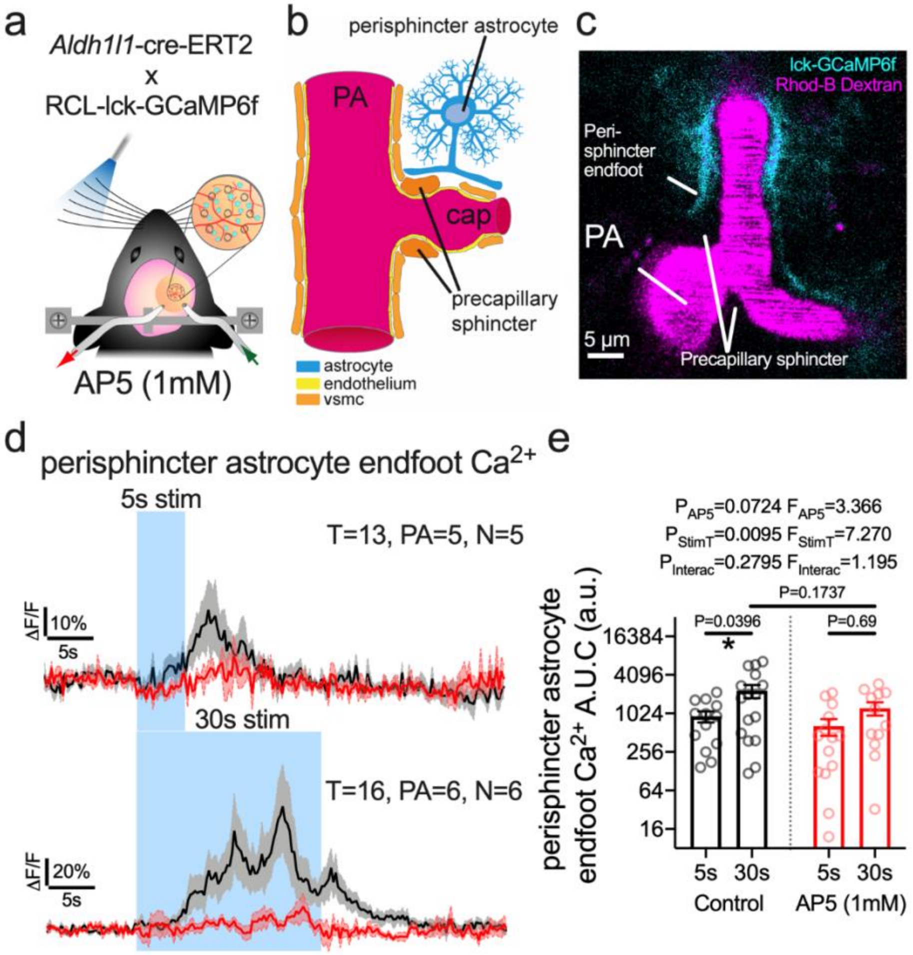


**<u>Supplemental Figure 8</u>: Peri-sphincter astrocyte $Ca^{2+}$ in response to 5sec and 30sec whisker stimulation**. **a**) Cartoon of *in vivo* experimental setup using membrane tethered GCaMP6f in astrocytes. **b**) Cartoon depicting astrocyte of interest (blue), adjacent to a pre-capillary sphincter. **c**) 2-photon image of a penetrating arteriole (PA) (magenta) and a narrowing at the first branch off the penetrator where mural sphincter cells are located. Surrounding astrocytes expressing membrane targeted lck-GCaMP6f are shown. **d**) Average time series trace data of astrocyte endfoot $Ca^{2+}$ in pre-drug control (black) and in the presence of AP5 (red) surrounding an arteriole sphincter in response to 5sec or 30sec whisker stimulation. **e**) *S*ummary data of AUC.

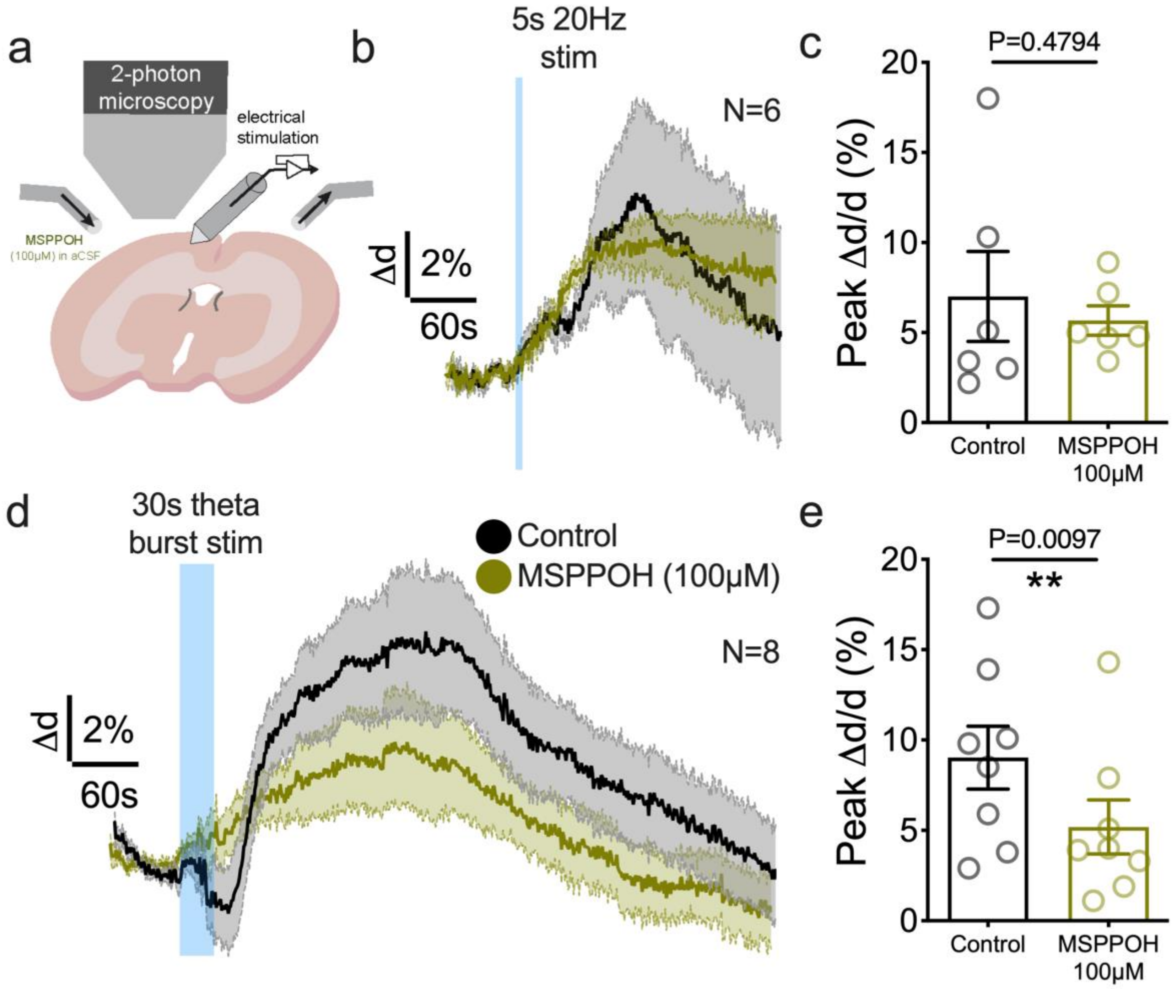


**<u>Supplemental Figure 9</u>: Epoxygenase inhibition with MSPPOH reduces arteriole dilation to 30sec high frequency afferent stimulation but not to 5sec**. **a**) Cartoon of experimental brain slice setup with electrical afferent stimulation. **b**) Average traces of evoked arteriole dilation to 5sec stim in pre-drug control (black) and in the presence of MSPPOH (green). **c**) Summary data for 5sec stim, showing no effect. **d**) Average traces of evoked arteriole dilation to 30sec electrical stim in pre-drug control (black) and in the presence of MSPPOH (green). **e**) Summary data for 30sec stim, showing a significant reduction.

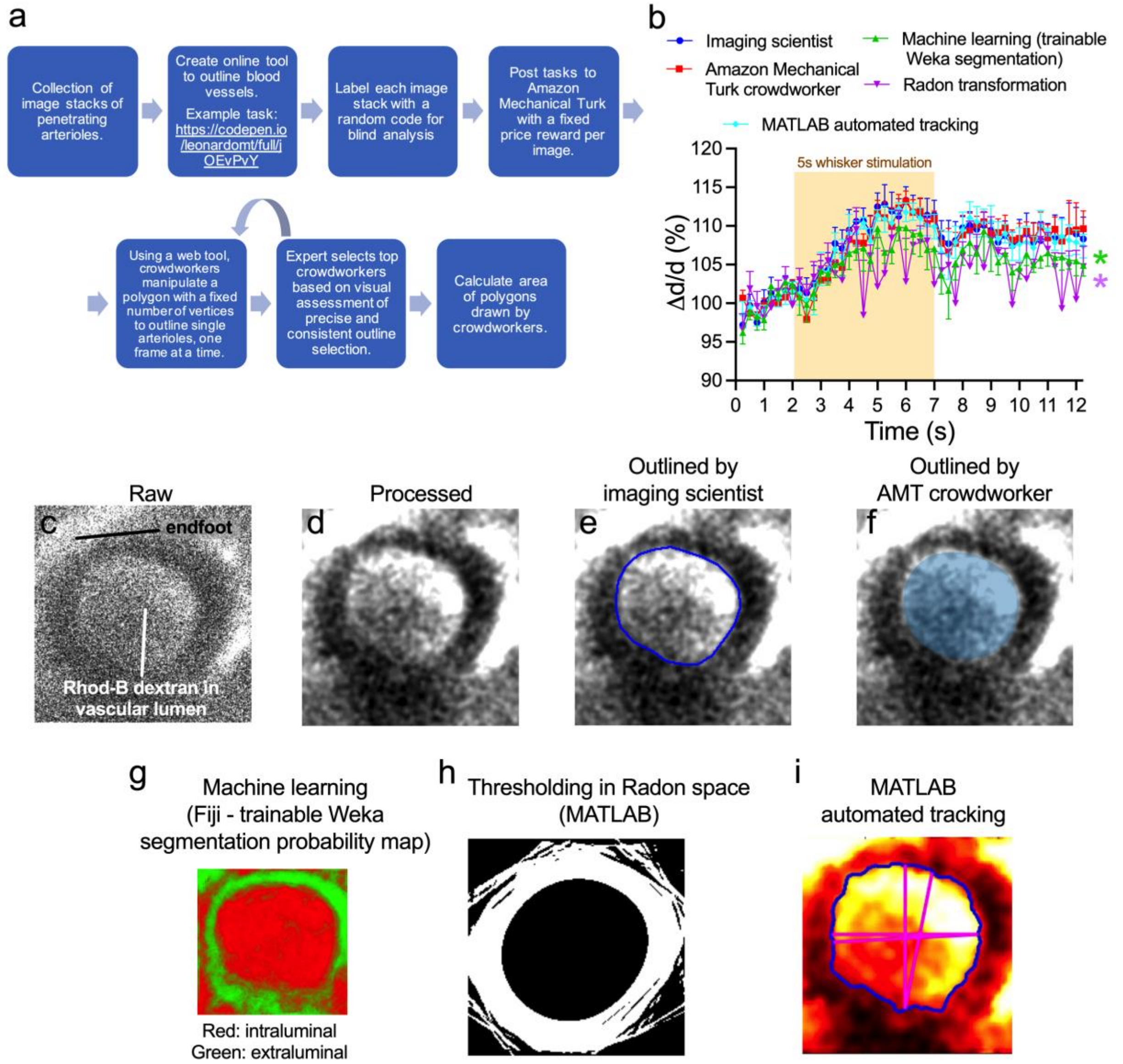


**Supplemental Figure 10: Validation of crowd sourced analysis of arteriole diameter changes**. **a**) Workflow of analysis using Amazon Turk with validation by imaging scientist. **b**) Analysis of arteriole diameter changes by 'trained' crowd-workers sourced via Amazon Turk performed equally well as a trained imaging scientist. Both these analyses outperformed an ImageJ machine learning tool called WEKA as well as implementing a radon transform of the data (PMID 24736890). **c-i**) representative images of arteriole lumen pre-processing ***(c,d)*** followed by the identification of the arteriole lumen by either an imaging scientist ***(e)*** or a crowd-worker ***(f)***,

WEKA segmentation ***(g)*** thresholding in Radon Space ***(h)*** MATLAB automated tracking based on Thirion's DEMONS algorithm*.*
